# *Salmonella* Typhimurium Vitamin B12-dependent Methionine metabolism regulates *C. elegans* Development

**DOI:** 10.64898/2026.01.27.702016

**Authors:** Chamjailiu Daimai, Vidya Devi Negi

## Abstract

Host-microbe interactions extend beyond infection, involving microbial metabolites such as short-chain fatty acids, vitamins, and other bioactive compounds that influence host physiology and development. Using *Caenorhabditis elegans,* an excellent model organism for host-microbe interactions, we demonstrate that *Salmonella* Typhimurium significantly accelerates *C. elegans* growth through a vitamin B12-dependent mechanism affecting methionine metabolism. We measured the reproductive maturity of *C. elegans* via embryo and oocyte production in response to various *Salmonella* virulence factor mutants and *Salmonella* strains deficient in cobalamin, folate, and methionine synthesis pathways. We found that the *Salmonella cbiB* and *metH* genes are essential for rapid development in *C. elegans*. Moreover, the *C. elegans metr-1* mutant, which lacks functional methionine synthase, did not exhibit increased reproductive maturity when fed on *S.* Typhimurium or *E. coli* OP50 supplemented with vitamin B12. However, methionine supplementation rescued this phenotype, indicating that *C. elegans* obtains vitamin B12 from *S.* Typhimurium, enhancing the methionine/S-adenosylmethionine (MET/SAM) cycle. We further observed that the *C. elegans sams-1* mutant, deficient in SAM synthase, showed growth defects and reduced oocyte production when fed on *Salmonella* or OP50 supplemented with vitamin B12 or methionine, respectively. Thus, SAM emerges as a key link between bacterial vitamin B12 metabolism and host reproduction.

Additionally, *Salmonella* increases vitellogenin-2 expression, reduces glutathione S-transferase activity, and decreases mitochondrial function. However, Δ*cbiB* and Δ*metH* reversed the beneficial effects of vitamin B12-dependent methionine on vitellogenesis and oxidative stress, suggesting that *Salmonella* promotes vitellogenesis and reduces oxidative stress through vitamin B12-dependent mitochondrial regulation, linking microbial metabolism to host development

**IMPORTANCE:** Understanding the interactions between hosts and pathogens is essential for improving host health. *C. elegans* obtains micronutrients from diet, gut microbiota, and supplements, but the contribution of microbiota in nutrient absorption is not fully understood. Bacterial metabolism may influence micronutrient status and pathogenicity, producing positive and adverse health effects on the host. This study demonstrates that pathogenic *Salmonella* can serve as a nutritional source for *C. elegans* when the two organisms coexist in the same environment. The findings suggest that, rather than being solely detrimental, *Salmonella* can alter the dietary habits of *C. elegans*, highlighting a complex interaction between the pathogen and its host. This relationship may provide insights into the ecological dynamics of microbial communities and their influence on host physiology and development.

## INTRODUCTION

Host-pathogen interactions are complex and involve multiple regulatory mechanisms. *Caenorhabditis elegans (C. elegans)* encounters various bacteria in its natural habitat, including pathogenic ones (1, 2). The interplay between a host response and pathogen virulence is intricate (3, 4). Host metabolism shifts to secure essential resources during infection, driven by immune activation and pathogen strategies (5). Pathogens compete with the host for nutrients, altering infection outcomes (6). Consequently, host metabolic pathways adapt to stress and reallocate nutrients to meet physiological needs during infection (7). *C. elegans* has emerged as a valuable model for studying immunometabolism (8). *C. elegans* has a three-day lifecycle progressing through embryogenesis, four larval stages (L1-L4), and adulthood (9, 10). Host-pathogen interactions influence reproductive physiology, but few studies have examined the impacts of pathogens on reproduction (11, 12). Using *C. elegans*, we investigated how *Salmonella* Typhimurium (*S.* Typhimurium) affects host reproduction. *S*. Typhimurium also causes major gastrointestinal and systemic diseases in humans and animals (13). The pathogenesis of *Salmonella* is primarily regulated by the virulence plasmid (pSLT), particularly through the *spv* operon, secretion systems, and *Salmonella* Pathogenicity Islands (SPIs) (14, 15). *Salmonella* infection accelerates larval development, reduces lifespan, and induces developmental changes linked to host pathways and *Salmonella* virulence factors, highlighting how bacterial pathogens can modulate *C. elegans* development (16).

*C. elegans* obtains B vitamins, folate, vitamin B12, vitamin B6, and iron through dietary intake from bacteria and bacterial metabolism in the gut microbiota (17, 18). Vitamin B12 is a key micronutrient produced only by certain prokaryotes, but is needed by various animals, including *C. elegans* (19, 20). Vitamin B12 is an essential cofactor in two key metabolic pathways. It aids methylmalonyl-CoA mutase (MUT, MMCM-1 in *C. elegans*) in mitochondrial propionate metabolism. It supports methionine synthase (MS, METR-1 in *C. elegans*) in one-carbon metabolism, including the methionine and folate cycles (21, 22). Vitamin B12 deficiency is associated with diverse clinical pathologies, including anaemia, neuropathy, and congenital disabilities (23). In *C. elegans,* it leads to infertility, slower growth rate, and reduced lifespan (24). *Escherichia coli (E. coli)* cannot synthesize vitamin B12 de novo and solely produces vitamin B12 through the salvage pathway. Hence, the levels of vitamin B12 in *E. coli* are significantly lower compared to species such as *Salmonella* and *Comamonas,* which can synthesize it de novo. The methionine biosynthetic pathway, highly conserved across prokaryotes, is absent in vertebrates, making it a plausible antimicrobial target (25). However, de novo methionine biosynthesis is essential in *S.* Typhimurium. Methionine is crucial for protein synthesis and S-adenosylmethionine (SAM) production, supporting methylation and animal reproductive output (26, 27). It influences the expression of the vitellogenin gene, affecting yolk production and reproductive fitness. Consequently, the availability of methionine directly impacts vitellogenesis and embryo development in *C. elegans* and other species (28–30). In addition to yolk formation, vitellogenins in *C. elegans* also enhance the host’s defense against pathogens (31). Exposure to certain bacteria accelerates reproductive maturity without impacting total reproduction (32, 33). Notably, methionine metabolism plays a crucial role in balancing oxidative stress by maintaining mitochondrial integrity, providing cysteine for glutathione synthesis, and facilitating methylation reactions that regulate antioxidant defenses (34, 35). Optimal reproductive success depends on the coordinated activity of methionine metabolism, efficient antioxidant defense, and healthy mitochondrial function. Disruption in any component can lead to impaired fertility, defective oocyte provisioning, or increased embryonic mortality, thereby linking metabolic health to reproduction (36, 37).

## RESULTS

### *Caenorhabditis elegans* development alteration is independent of *Salmonella* virulence

Previously, the lab has shown how *S.* Typhimurium (WT-STM) infection of *C. elegans* leads to rapid development compared to *E. coli* OP50 (OP50) (16). In the current study, we exposed the *C. elegans* to WT-STM and several *Salmonella* mutants, including Δ*hilA,* Δ*avrA,* Δ*sptP,* Δ*sipD,* Δ*ssaV,* Δ*ygiM,* Δ*rspA,* Δ*sopB,* Δ*mgtC,* Δ*phoP*, and Δ*fepB,* to understand the involvement of major virulence factors regulating *C. elegans* development (Fig. 1A). Most of these mutants are avirulent or attenuated deletion strains, listed in Table 1 (38–45). *C. elegans* embryo production was quantified at 56 hours post-L1 stage to observe the developmental dynamics upon infection with these strains. Interestingly, the mutants showed accelerated development, with embryo production comparable to that of WT-STM (Fig. 1B). The number of oocytes produced was also quantified in day-1 adult *C. elegans.* A similar outcome was observed, with increased oocyte production during infection with the WT-STM and its avirulent mutant strains (Fig. 1C). This indicates that *C. elegans* exposed to WT-STM and its avirulent mutant strains attain early reproductive maturity, suggesting that these genes do not play a role in the enhanced development of *C. elegans*. We aimed to identify signaling pathways involved in WT-STM infection and its mutants’ mediated enhanced development by analyzing gene expression of Insulin/IGF-1 signaling, the RAS/MAPK pathway, and programmed cell death associated with oogenesis in *C. elegans* (46, 47). The pathway is initiated by the binding of LIN-3 (EGF-like ligand) to the LET-23 receptor tyrosine kinase (RTK) on the surface of vulva precursor cells (VPCs). A kinase cascade is activated with the downstream kinase MPK-1, which is activated in the pachytene and oocyte regions of the germline (48, 49). PAR-5 is a negative regulator of MPK-1 signaling, restricting its activity in the proximal region (50). In contrast, GLP-1/Notch signaling in the distal germline promotes germline stem cell (GSC) proliferation and prevents premature entry into meiosis (51–53). IGF-1 is involved in various processes, including growth, metabolism, and lifespan, and it is activated when insulin-like peptides bind to the DAF-2 receptor, initiating a signaling cascade (54). Additionally, CED-9 inhibits CED-4, and upon binding to EGL-1, it initiates programmed cell death by releasing CED-4, which then oligomerizes with the active caspase CED-3 to execute cell death (55, 56). However, the mRNA expression levels of these major target genes, such as *lin-3, let-23, mpk-1, glp-1, par-5, ced-9, ced-4, ced-3, daf-2,* and *ins-7*, suggest that they were not responsible for the rapid reproductive maturity observed during *Salmonella* infection (Fig. S1A-C). There was no increase in the life span of *C. elegans* exposed to various major SPI-1 effector mutants (Fig. S1D) as compared to WT-STM. These observations suggest that S. Typhimurium’s virulence factors do not contribute to the accelerated development of *C. elegans*. Instead, the presence of these bacteria in the diet may significantly affect the host’s development rather than its virulence.

**Fig. 1.**
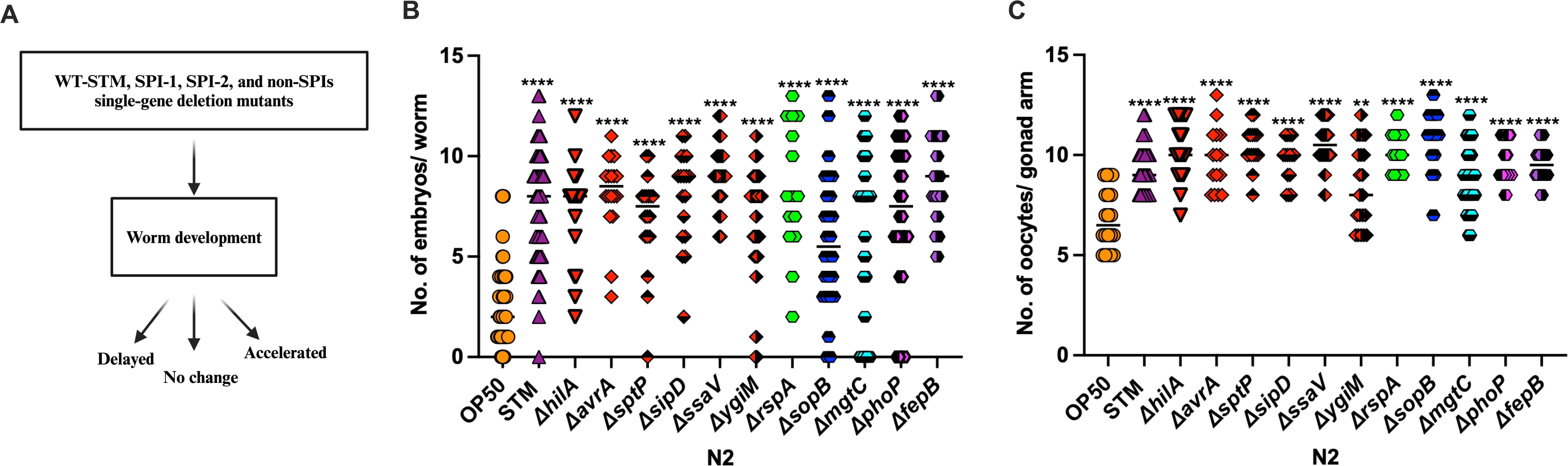
*Caenorhabditis elegans* development alteration is independent of *Salmonella* virulence. (A) Schematic representation of the screens conducted on the worm developmental rate. (B) Represents the quantification of embryos in N2 worms post 56 hours L1 stage and (C) Oocyte production post 72 hours L1 stage, which increases in WT-STM*, ΔhilA*, *ΔavrA*, *ΔsptP*, *ΔsipD*, *ΔssaV*, *ΔygiM*, *ΔrspA*, *ΔsopB*, *ΔmgtC*, *ΔphoP*, and *ΔfepB*-exposed animals. An ordinary one-way ANOVA, post-hoc Tukey HSD test was performed. *\*\*\*\*P<0.0001*; *\*\*\*P < 0.001; **P < 0.01; * P < 0.05*.

**Table 1.**
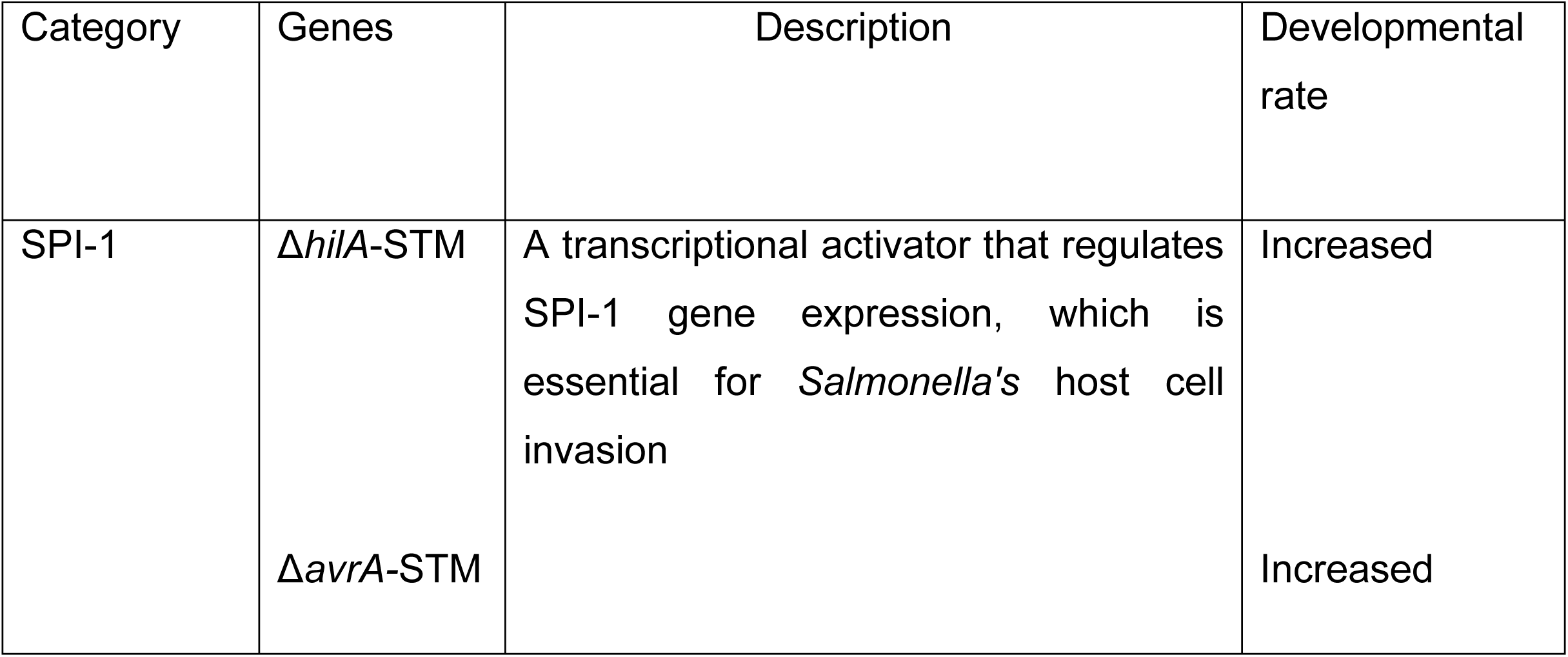
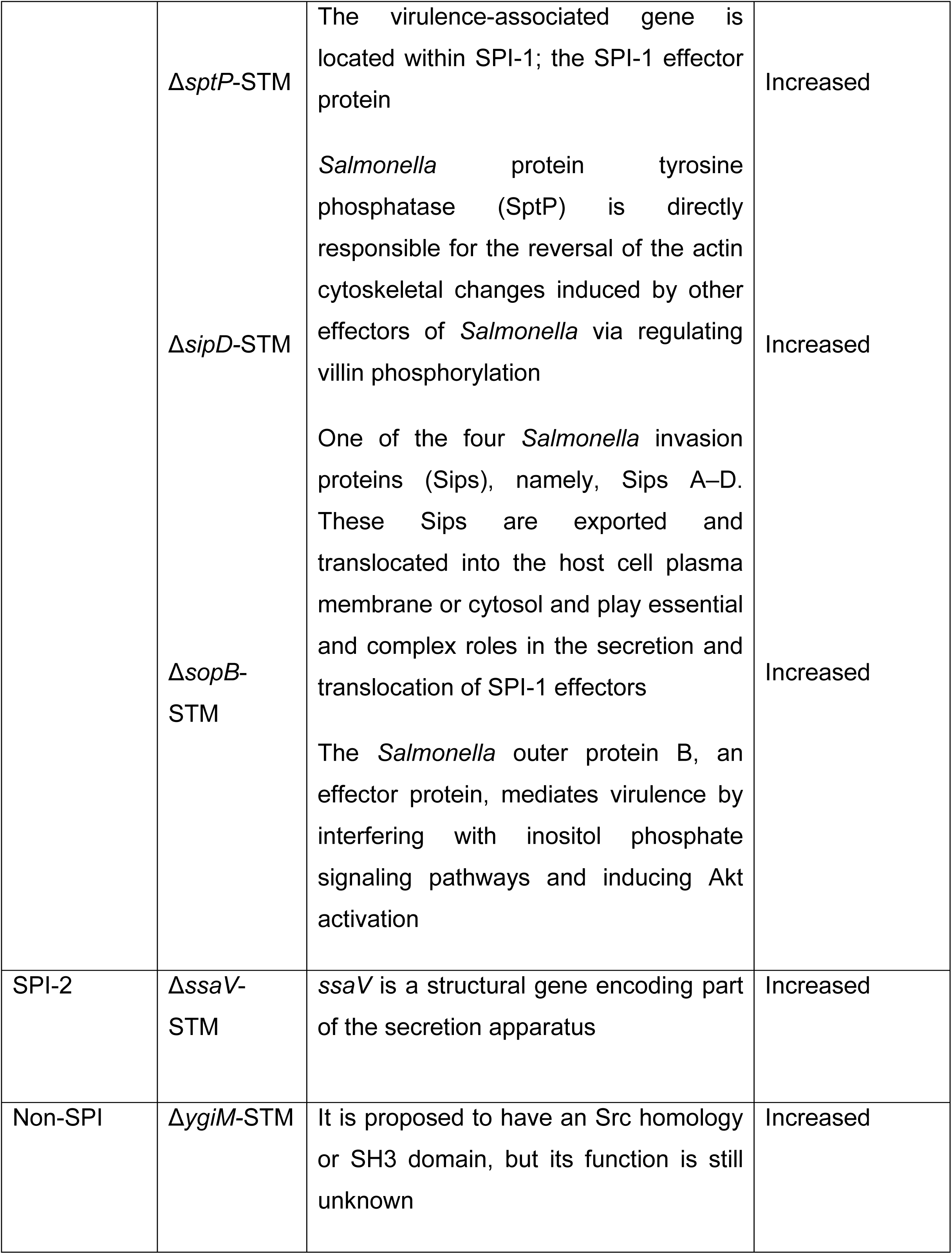
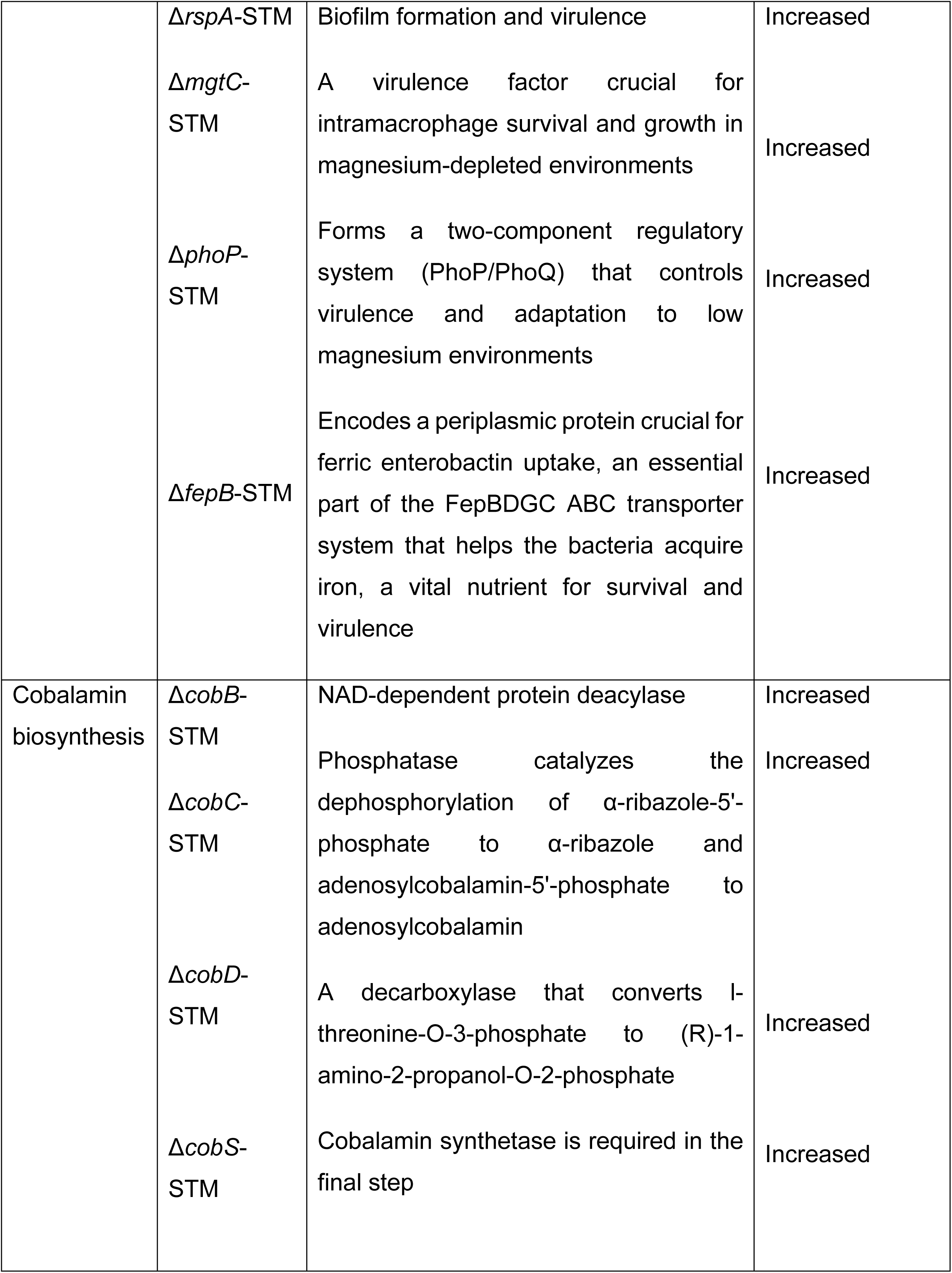
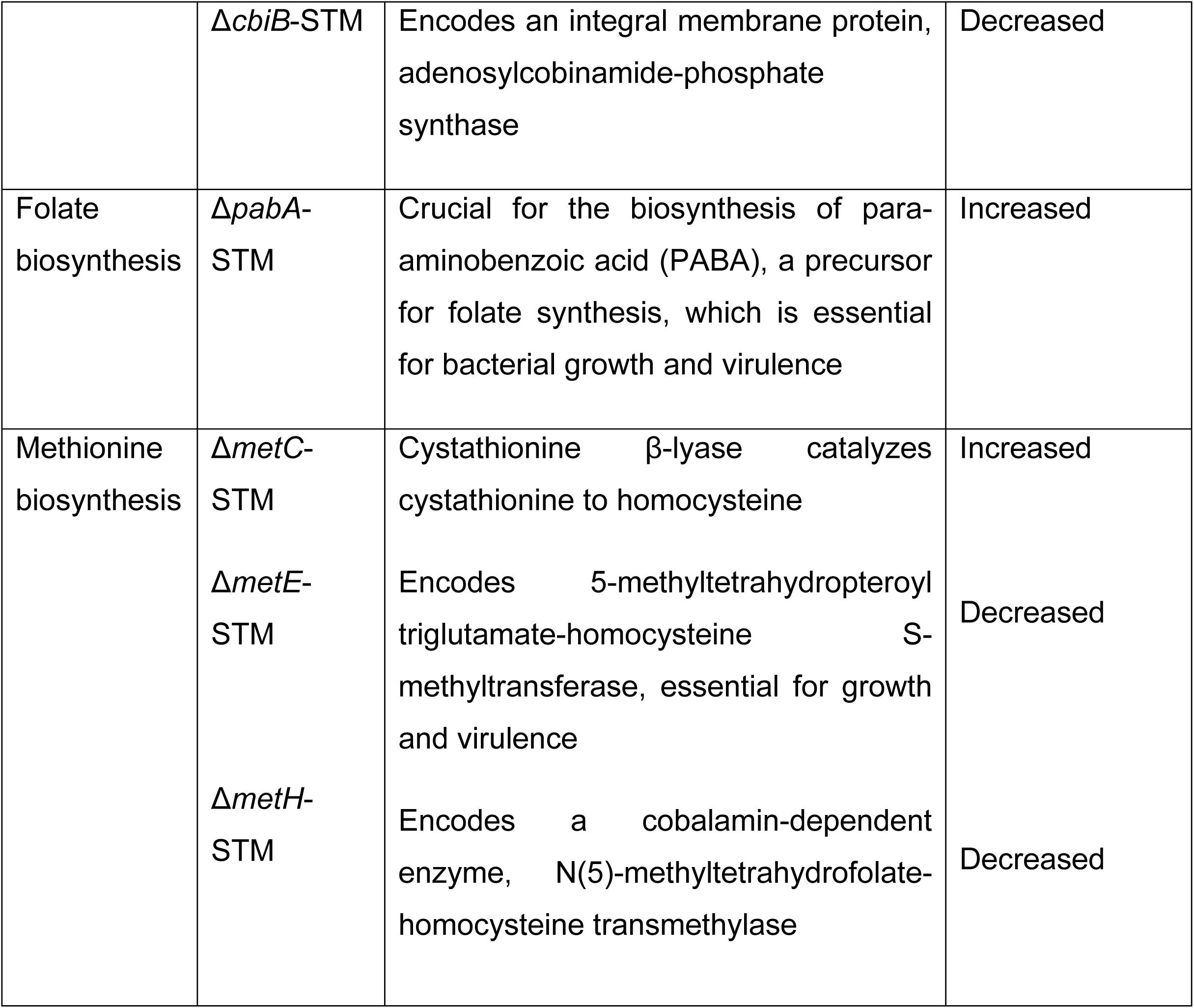
SPI-1, SPI-2, non-SPI-1, cobalamin, folate, and methionine biosynthesis single-gene deletion *Salmonella* mutants on developmental rate.

### *Salmonella* Typhimurium, a vitamin B12-rich diet, accelerates development and oocyte production in *C. elegans*

*Comamonas aquatica* DA1877 is known to accelerate *C. elegans* development (57) via the production of vitamin B12, a crucial cofactor in one-carbon metabolism. Animals acquire vitamin B12 either through a symbiotic relationship with gut bacteria or through their diet. Although *E. coli* can import vitamin B12 from standard growth media, the concentrations are significantly lower than those found in vitamin B12-producing bacteria, such as *Comamonas aquatica (C. aquatica)* (21). Unlike *Salmonella* and *Comamonas,* which can synthesize cobalamin de novo, *E. coli* lacks this capability and relies solely on the salvage pathway, resulting in a significantly lower intracellular vitamin B12 level (58). To investigate the role of bacterial metabolism in host reproduction, we quantified the number of progenies produced by the *C. elegans* exposed to *E. coli* OP50, WT-STM, Δ*hilA*, and *E. coli* OP50 supplemented with vitamin B12. The Δ*hilA* strain, a SPI-1 master regulator mutant, was used as a control to exclude the pathogenic effects on host reproduction. The number of progenies produced by the worm exposed to WT-STM, Δ*hilA*, and *E. coli* OP50 supplemented with vitamin B12 were increased significantly at the first two days, however by day 3, the number decreases and were comparable to the OP50, likely due to the depletion of the self-fertilized sperm as single hermaphrodites are being transferred to the assayed plates (Fig. 2A). We next examined the oocytes production in day-1 adult *C. elegans* using the OD95, a *pie-1p::mCherry::his-58 + unc-119(+)* reporter strain and observed a significant increase in production of oocytes in WT-STM, Δ*hilA*, and *E. coli* OP50 supplemented with vitamin B12 exposed *C. elegans* (Fig. 2B and C). This suggested that vitamin B12, rather than the bacteria’s virulence factor, contributed to the enhanced development observed. Additionally, to directly investigate the role of One-Carbon Metabolism (OCM), we created *Salmonella* mutants that are deficient in folate (Δ*pabA*), cobalamin (Δ*cobB*, Δ*cobC,* Δ*cobD,* Δ*cobS,* and Δ*cbiB*), and methionine (Δ*metC,* Δ*metE*, and Δ*metH*) synthesis. In *S. enterica*, folate-dependent OCM is essential for purines, thymidylate, and methionine biosynthesis, as well as for redox balance and translation regulation (59). We also used the VL749 strain, a transgenic worm that carries an *acdh-1::GFP* reporter, allowing it to detect the presence of vitamin B12 in the environment (60). A reduction in GFP fluorescence indicates the presence of vitamin B12. The worms were exposed to *E. coli* OP50, WT-STM, and *E. coli* OP50 supplemented with vitamin B12, as well as *Salmonella* mutants Δ*cobB,* Δ*cobC,* Δ*cobD*, Δ*cobS*, Δ*cbiB,* and Δ*pabA*. Our findings showed that ACDH-1::GFP expression was reduced significantly in the worm fed with *Salmonella* compared to those fed with *E. coli* OP50 and was almost absent in *E. coli* OP50 supplemented with vitamin B12 (Fig. 2D). We observed a restoration of GFP levels in animals fed with the cobalamin mutants, such as Δ*cobC,* Δ*cobD,* Δ*cobS*, and Δ*cbiB*. The GFP expression in the *acdh-1::GFP* worm indicates the role of dietary vitamin B12, as the *acdh-1* gene is a highly sensitive marker for vitamin B12 deficiency (20, 21). Importantly, no cobalamin, folate, or methionine synthesis mutants of *Salmonella* showed growth defects in LB media (Fig. S2A and S2B). Thus, these findings together suggest that *S.* Typhimurium accelerates development in *C. elegans* predominantly through dietary vitamin B12, rather than through mechanisms associated with infection or virulence.

**Fig. 2.**
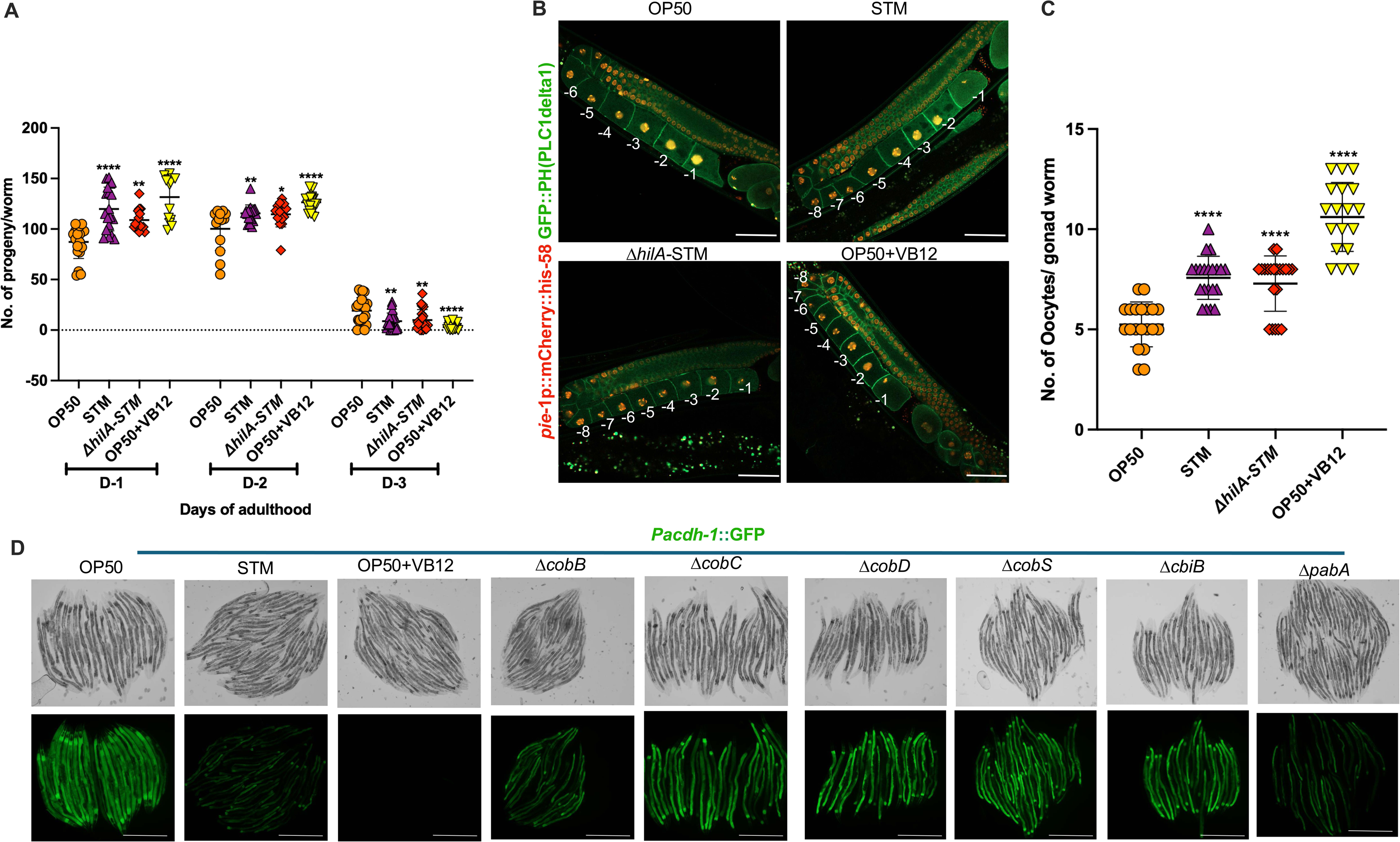
*S.* Typhimurium, a vitamin B12-rich diet accelerates reproductive maturity *in C. elegans.* (A) Wildtype *S.* Typhimurium, *ΔhilA-*STM, and vitamin B12-supplemented with *E. coli* OP50 increase the number of progeny within the first 2 days. (B) Representative confocal microscopy images of the gonad, using the OD95 (*ltIs37 [pie-1p::mCherry::his-58 + unc-119(+)] IV. ltIs38 [pie-1p::GFP::PH(PLC1delta1) + unc-119(+)])* transgenic strain, showing increased oocyte production in WT-STM, *ΔhilA-*STM, and OP50 supplementation with vitamin B12-fed day 1 worms. The numbers -1 and -2 indicate the most proximal mature oocytes. (C) Graphical data illustrating increased oocyte production in WT-STM, *ΔhilA-*STM, and OP50 supplementation with vitamin B12-fed worms. (D) Representative fluorescence images of *acdh-1p::GFP* animals grown on *E. coli* OP50, wild-type *Salmonella* Typhimurium (WT-STM), cobalamin (Δ*cobB,* Δ*cobC,* Δ*cobD*, Δ*cobS*, Δ*cbiB*), and folate synthesis (Δ*pabA) Salmonella* mutants, and *E. coli* OP50 supplemented with Vitamin B12, where GFP signals were imaged. An ordinary one-way ANOVA, post-hoc Tukey HSD test was performed. *\*\*\*\*P<0.0001*; *\*\*\*P < 0.001; **P < 0.01; * P < 0.05.* Scale bar = 100 μm.

### *Salmonella* cobalamin (*cbiB*) and methionine (*metH*) synthesis mutants reduce the developmental rate of *C. elegans*

Considering the essential and critical role of one-carbon metabolism in *Salmonella* in synthesizing nucleotides, amino acids, and cofactors, we were interested in understanding whether disrupting this pathway could mitigate the accelerated reproductive maturity of *C. elegans*. Aged-synchronized L1 larvae were fed on *E. coli* OP50, WT-STM, *E. coli* OP50 supplemented with vitamin B12, Δ*cobB,* Δ*cobC,* Δ*cobD,* Δ*cobS,* Δ*cbiB*, *E. coli* OP50 supplemented with methionine, Δ*metC,* Δ*metE*, and Δ*metH*. Embryo production was quantified at 56 hours post-L1 stage to assess developmental acceleration. Interestingly, the Δ*cbiB* strain exhibited reduced embryo production, but this was restored by vitamin B12 supplementation (Fig. 3A). Similarly, *C. elegans* fed with Δ*cobB,* Δ*cobC,* Δ*cobD,* Δ*cobS*, and Δ*cbiB* produced fewer oocytes. This phenotype was reversed by vitamin B12 supplementation (Fig. 3B). The animals fed with Δ*metE* and Δ*metH* showed reduced embryo production, which was reversed with methionine supplementation (Fig. 3C). Consistently, the worms fed with Δ*metC,* Δ*metE,* and Δ*metH* produced fewer oocytes. Phenotype was reversed with methionine supplementation (Fig. 3D). Together, these findings demonstrate that the *Salmonella* one-carbon metabolism pathway is crucial in regulating the reproductive maturity of *C. elegans*.

**Fig. 3.**
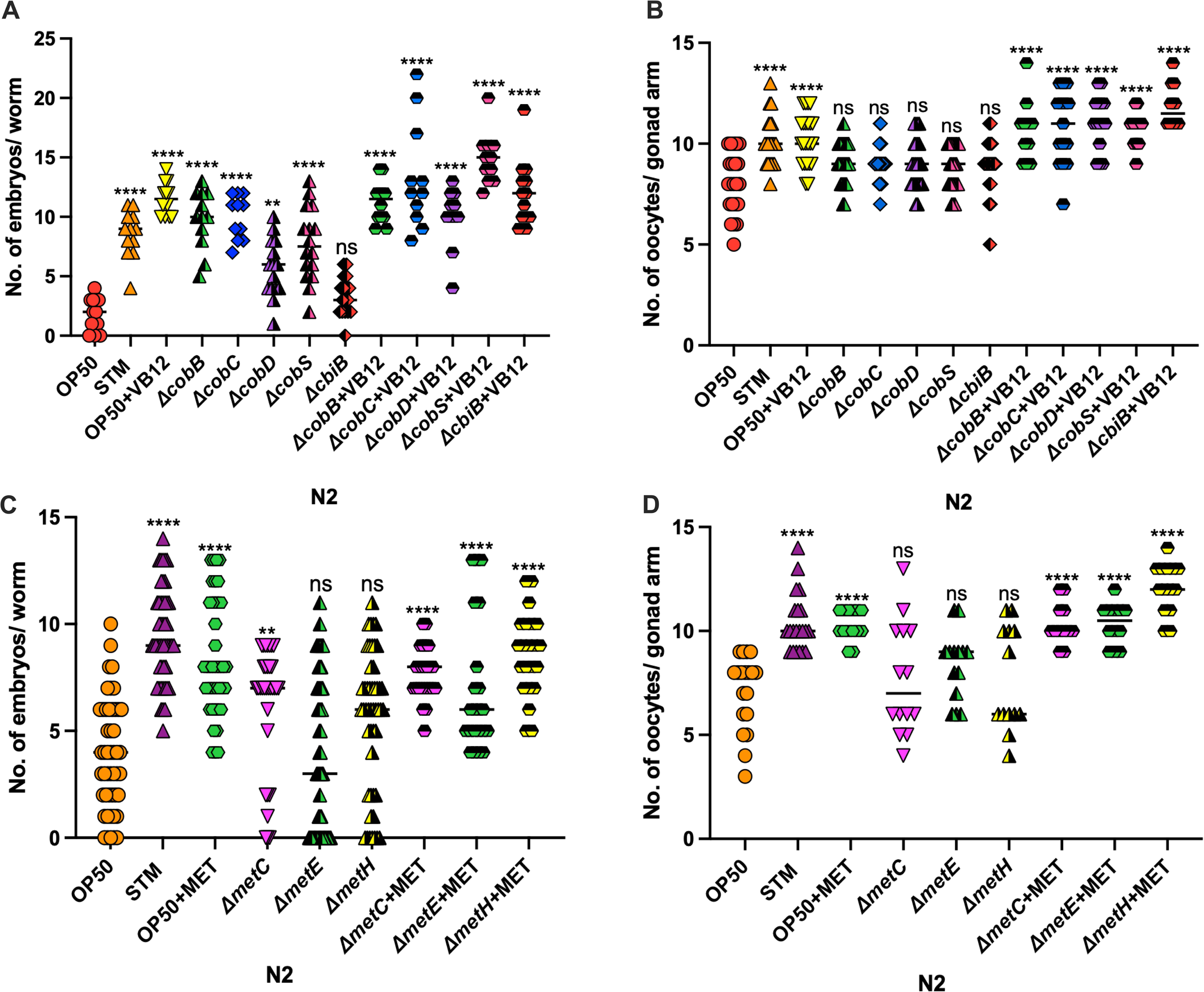
*Salmonella* mutants deficient in cobalamin (*cbiB*) and methionine (*metH*) synthesis reduce the developmental rate of *C. elegans.* (A) Represents the quantification of embryos in N2 worms post 56 hours L1 stage, fed on *E. coli* OP50, WT-STM, *E. coli* OP50 supplemented with vitamin B12, Δ*cobB,* Δ*cobC,* Δ*cobD,* Δ*cobS,* Δ*cbiB*, vitamin B12 supplemented with Δ*cobB,* Δ*cobC,* Δ*cobD,* Δ*cobS,* and Δ*cbiB,* respectively. *cbiB Salmonella* had lower embryo production at 56 hours post-L1, and supplementation with vitamin B12 increased the embryo number. (B) Represents the quantification of oocyte production in N2 worms post 72 hours L1 stage on *E. coli* OP50, WT-STM, *E. coli* OP50 supplemented with vitamin B12, Δ*cobB,* Δ*cobC,* Δ*cobD,* Δ*cobS,* Δ*cbiB*, vitamin B12 supplemented with Δ*cobB,* Δ*cobC,* Δ*cobD,* Δ*cobS,* and Δ*cbiB,* respectively. The *cbiB* mutant reduces the oocyte production, with the reversal observed in vitamin B12 supplementation. (C) Represents the quantification of embryos in N2 worms post 56 hours L1 stage, fed on *E. coli* OP50, WT-STM, *E. coli* OP50 supplemented with methionine, Δ*metC,* Δ*metE,* Δ*metH,* vitamin B12 supplemented with Δ*metC,* Δ*metE,* and Δ*metH*, respectively. *metE* and *metH* mutants decreased embryo and (D) oocyte production, with the reversal observed in methionine supplementation, respectively. An ordinary one-way ANOVA, post-hoc Tukey HSD test was performed. *\*\*\*\*P<0.0001*; *\*\*\*P < 0.001; **P < 0.01; * P < 0.05*.

### *Salmonella* accelerates development through vitamin B12-dependent metabolism in *C. elegans*

The *C. elegans* one-carbon metabolism pathway, primarily involving folate and methionine cycles, is essential for nucleotide synthesis, methylation reactions (61), DNA repair, and gene expression regulation (62). While direct interactions between *Salmonella’s* one-carbon metabolism and the host’s metabolic pathways have not been extensively studied, it’s plausible that the pathogen’s metabolic activities could influence these host pathways. Thus, to further investigate whether *Salmonella’s* vitamin B12-dependent one-carbon metabolism influences host development, we exposed the *metr-1, sams-1*, and *mmcm-1* mutant strains of the *C. elegans* to the above exposure conditions and assessed embryo and oocyte production. In *metr-1* mutants, the vitamin B12-dependent methionine synthesis pathway is impaired, and vitamin B12 supplementation did not rescue the defects (20, 63). Thus, the mutant continued to exhibit methionine-deficiency phenotypes, including developmental delays and growth defects (21, 61). Methionine is an essential amino acid required to make S-adenosylmethionine (SAM), a key methyl donor in many biochemical reactions (64, 65). However, supplementing *E. coli* OP50 with exogenous methionine bypasses the defective pathway in *metr-1* mutants (Fig. 4B and 4F), possibly by restoring the methionine and SAM pools and rescuing both growth and reproductive defects in the worm. Δ*cbiB* and Δ*metH*, which are cobalamin and methionine biosynthesis mutants, respectively, caused a delay in development with comparable embryo production *to E. coli* OP50 and a defect in oocyte production in *metr-1* animals (Fig. 4B and 4F). This suggests that methionine availability from the bacterial diet is critical for host rescue. *Salmonella* produces more methionine due to high vitamin B12 levels; *metr-1* mutants can salvage methionine from their bacterial diet to rescue their development. However, WT-STM infection could not rescue the *metr-1* animals’ growth and developmental defects phenotype, indicating that *metr-1* cannot utilize the bacterial vitamin B12 to synthesize methionine. Exogenous methionine supplementation in *Salmonella* rescued the *metr-1* developmental delay, resulting in a rapid developmental phenotype and increased oocyte production (Fig. 4C and 4G). This indicates that the *metr-1* worms cannot acquire vitamin B12 from the bacterial source to synthesize methionine from homocysteine.

**Fig. 4.**
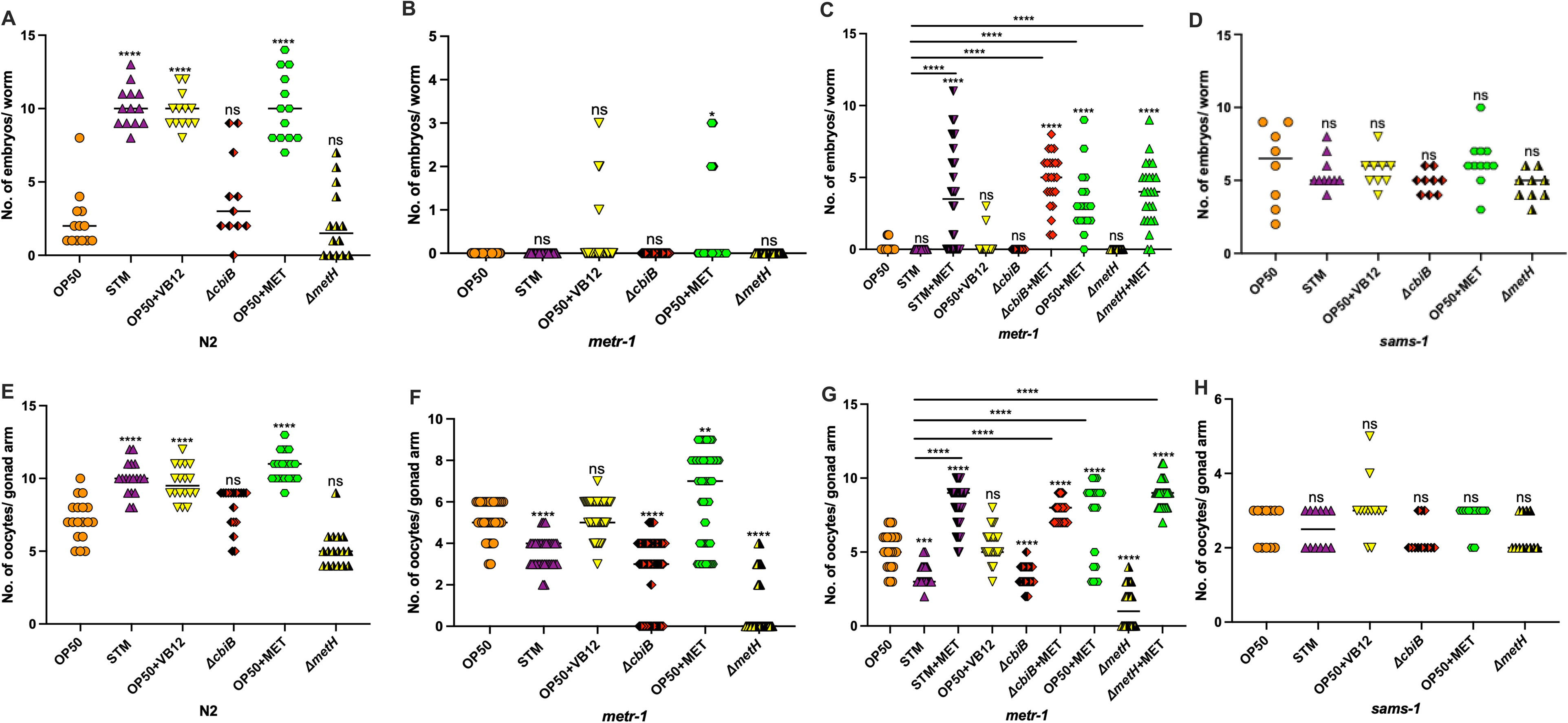
*Salmonella* Typhimurium accelerates development through vitamin B12-dependent metabolism in *C. elegans.* Represents the quantification of embryos in (A) N2, (B) *metr-1*, and (D) *sams-1* on *E. coli* OP50, WT-STM, *E. coli* OP50 supplemented with vitamin B12 and methionine, respectively, Δ*cbiB,* and Δ*metH*. (C) Represents the quantification of embryos in *metr-1* on *E. coli* OP50, WT-STM, *E. coli* OP50 supplemented with vitamin B12 and methionine, respectively, Δ*cbiB,* and Δ*metH,* methionine supplementation with WT-STM, Δ*cbiB,* and Δ*metH,* respectively. (E), (F) and (H) Represents the quantification of oocyte production in N2, *metr-1,* and *sams-1* animals on the indicated bacterial conditions as mentioned earlier. (G) Represents the quantification of oocytes in *metr-1* on *E. coli* OP50, WT-STM, *E. coli* OP50 supplemented with vitamin B12 and methionine, respectively, Δ*cbiB,* and Δ*metH,* methionine supplementation with WT-STM, Δ*cbiB,* and Δ*metH,* respectively. An ordinary one-way ANOVA, post-hoc Tukey HSD test was performed. *\*\*\*\*P<0.0001*; *\*\*\*P < 0.001; **P < 0.01; * P < 0.05*.

The *sams-1* mutant displayed growth defects and developmental anomalies (22, 66). Hence, we quantified the embryo production at 72 hours post-L1 stage and oocyte number at 96 hours post-L1 stage. Similarly, we found reduced embryo and oocyte production when exposed to different bacterial diets, including *E. coli* OP50, WT-STM, *E. coli* OP50 supplemented with vitamin B12, or methionine, resulting in growth defects (Fig. 4D and H). The data indicate that WT-STM, a facultative anaerobe, produces vitamin B12 in the gut of *C. elegans*, which the host utilizes to synthesize methionine, a precursor of SAM. The *mmcm-1* mutant, defective in the vitamin B12-dependent propionate breakdown pathway, shows no growth defects with these various bacterial diets (Fig. S3A and B). This indicates that methionine synthesis via vitamin B12 is essential for the rapid development and reproductive maturity of *C. elegans*. These results suggest that WT-STM produces vitamin B12 in the *C. elegans* gut, which the host uses to synthesize methionine and SAM. This *Salmonella* metabolic pathway accelerates and enhances early reproductive maturity in *C. elegans*.

### *Salmonella* enhances vitellogenin activity and mitigates oxidative stress, restoring vitellogenesis and oocyte development

Vitellogenins are yolk proteins synthesized in the adult intestine that supply nutrients to developing oocytes to support embryo development (67, 68). The *C. elegans* fed with WT-STM produced more oocytes (Fig. 2B). As yolk proteins are major components of oocytes, we further examined vitellogenin levels in worms fed on *Salmonella*. We assessed vitellogenin gene 2 (*vit-2*) expression using a *vit-2::GFP* reporter strain. We observed increased expression of VIT-2::GFP in both oocytes (Fig. 5A and B) and embryos (Fig. 5C and D) when *C. elegans* were fed on WT-STM or *E. coli* OP50 supplemented with vitamin B12 or methionine, respectively. In contrast, a comparable VIT-2::GFP expression was observed in *C. elegans* fed on Δ*cbiB* and *ΔmetH* and *E. coli* OP50. This suggests that a vitamin B12-rich diet of WT-STM enhances vitellogenesis and developmental growth by modulating methionine metabolism.

**Fig. 5.**
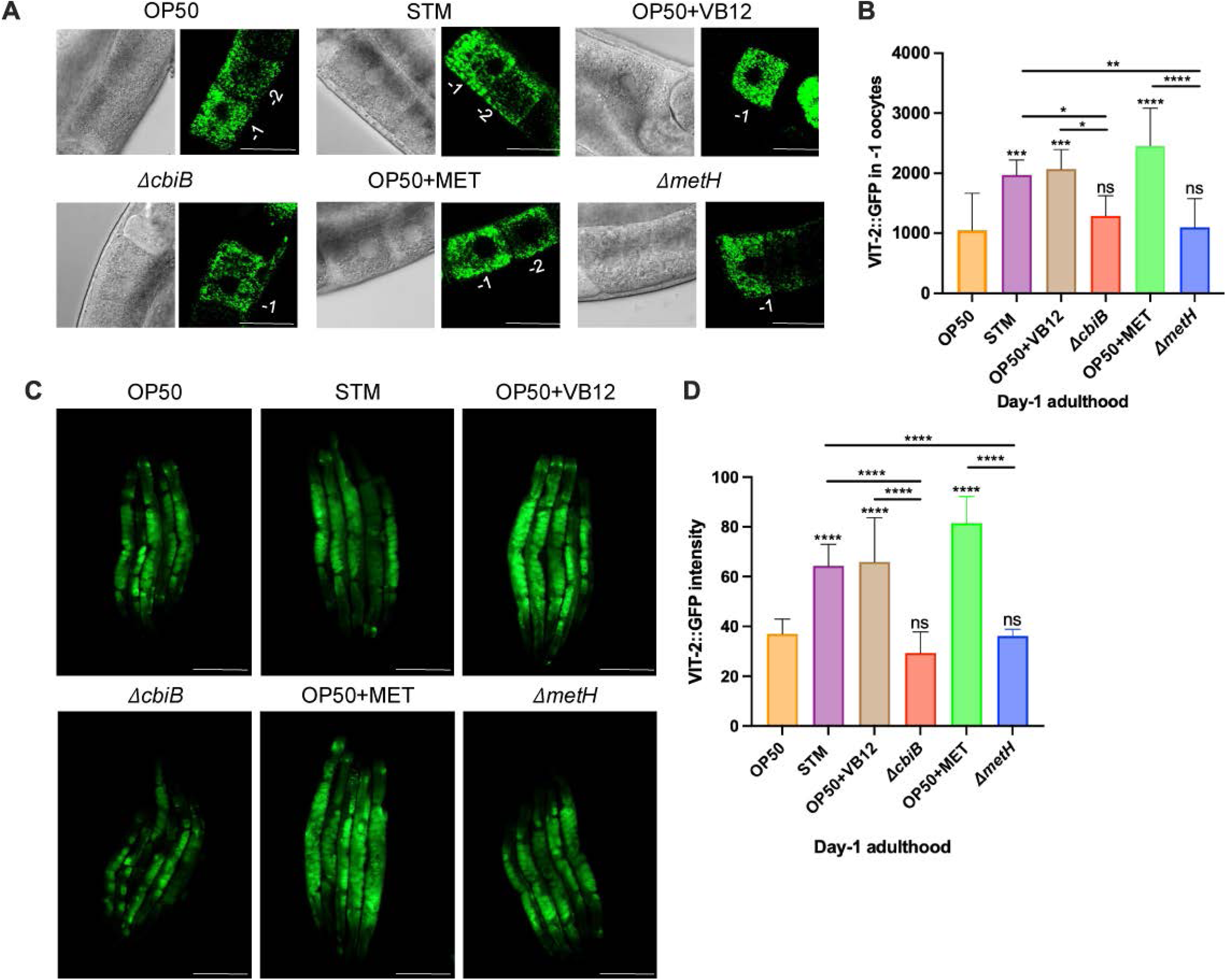
*Salmonella* Typhimurium increases vitellogenin activity, restoring vitellogenesis and oocyte development. (A) The representative confocal microscopy images of the vitellogenin-2 expression, using *vit-2::gfp* transgenic worms in the primary oocytes of day-1 adults fed on *E. coli* OP50, wildtype *Salmonella,* Δ*cbiB,* Δ*metH, E. coli* OP50 supplemented with either vitamin B12 or methionine, respectively. (C) The representative images and (D) Quantification of vitellogenin-2 expression in the embryos of day-1 adults under the indicated bacterial conditions. An ordinary one-way ANOVA, post-hoc Tukey HSD test was performed. *\*\*\*\*P<0.0001*; *\*\*\*P < 0.001; **P < 0.01; * P < 0.05.* Scale bar = 100 μm.

Vitamin B12 has also been shown to exhibit antioxidant activity, which helps mitigate oxidative stress (69, 70). We observed increased VIT-2 expression and oocyte production in *C. elegans* fed on *Salmonella* driven by vitamin B12-dependent methionine metabolism. Reactive oxygen species (ROS) are cleared by antioxidant enzymes, such as GST-4, a glutathione S-transferase regulated by *skn-1* during oxidative stress response (71–73). Therefore, we examined GST-4 levels using the *gst-4*::*GFP* reporter strain, which was fed on WT-STM or *E. coli* OP50 supplemented with vitamin B12 or methionine, respectively. Consistent with the role of vitamin B12, we found reduced GST-4 expression in WT-STM and *E. coli* OP50 supplemented with vitamin B12 or methionine, respectively, while *E. coli* OP50, Δ*cbiB*, and Δ*metH* fed *C. elegans* displayed increased GST-4 activation (Fig. 6A and B). ROS levels were measured by the intensity of Dichlorofluorescein (DCF), which were significantly higher in worms fed diets of *E. coli* OP50, Δ*cbiB*, and Δ*metH* compared to those on the vitamin B12-rich diet of WT-STM, *E. coli* OP50 supplemented with B12, and methionine, respectively (Fig. 6C). These findings indicate that the vitamin B12-rich diet enhanced vitellogenin activity, which helped to reduce oxidative stress in *C. elegans* fed on WT-STM. To confirm that the effect depends on methionine metabolism, we assessed ROS levels in the *metr-1* mutant, which cannot acquire dietary vitamin B12, a cofactor required for methionine synthesis, leading to decreased reproductive maturity. This suggests reduced vitellogenesis and increased oxidative stress. However, ROS levels in *metr-1* worms on vitamin B12-rich diets were comparable to *E. coli* OP50, Δ*cbiB,* and Δ*metH* (Fig. 6D). Interestingly, *Salmonella*, however, is reported to induce oxidative stress in *C. elegans* when L4 nematodes are acutely exposed for 24 hours (74). However, in our experiment’s paradigm, feeding differed, as we exposed L1 larvae to *Salmonella* until they reached adulthood. This condition may result in reduced ROS because *C. elegans* consumes *Salmonella* as its diet, leading to increased expression of VIT-2 and indicating enhanced antioxidant activity, which consequently reduces ROS. We observed that the vitellogenin levels in the nematodes decreased after 24 hours of exposure to *Salmonella* Typhimurium. In contrast, VIT-2 expression increased upon supplementation with *E. coli* OP50, vitamin B12, and methionine (Fig. S4A and B). This finding was further validated using the *gst::4-GFP* reporter strain, which showed no significant change in GST-4 expression compared to *E. coli* OP50 but exhibited reduced expression during *E. coli* OP50 supplementation with vitamin B12, and methionine, respectively (Fig. S4C and D). The methionine cycle also contributes to cysteine synthesis, a precursor for glutathione (75, 76), an important antioxidant that lowers oxidative stress, which otherwise suppresses VIT-2 expression (77) and ultimately supports oocyte development. Methionine metabolism is a key regulator of lifespan in *C. elegans*. Methionine restriction extends lifespan, while effective cycling is vital for dietary and metabolic interventions that promote longevity (78). Its links to reproduction, stress response, and epigenetics make methionine an important factor in aging in nematodes (77, 79). We observed that WT-STM exposure caused early worm mortality, while *E. coli* OP50 supplemented with methionine and Δ*cbiB* exposed worms showed no significant survival difference compared to *E. coli* OP50. However, *E. coli* OP50 supplemented with vitamin B12 and Δ*metH* led to longer survival than the other bacterial diets (Fig. S4E). Methionine metabolism plays a crucial role in the growth and virulence of *S.* Typhimurium; mutants deficient in these pathways are less virulent (13). This suggests the role of metabolic and virulence competence in *S.* Typhimurium, highlighting the importance of these metabolites to both the pathogen and the host (80, 81).

**Fig. 6.**
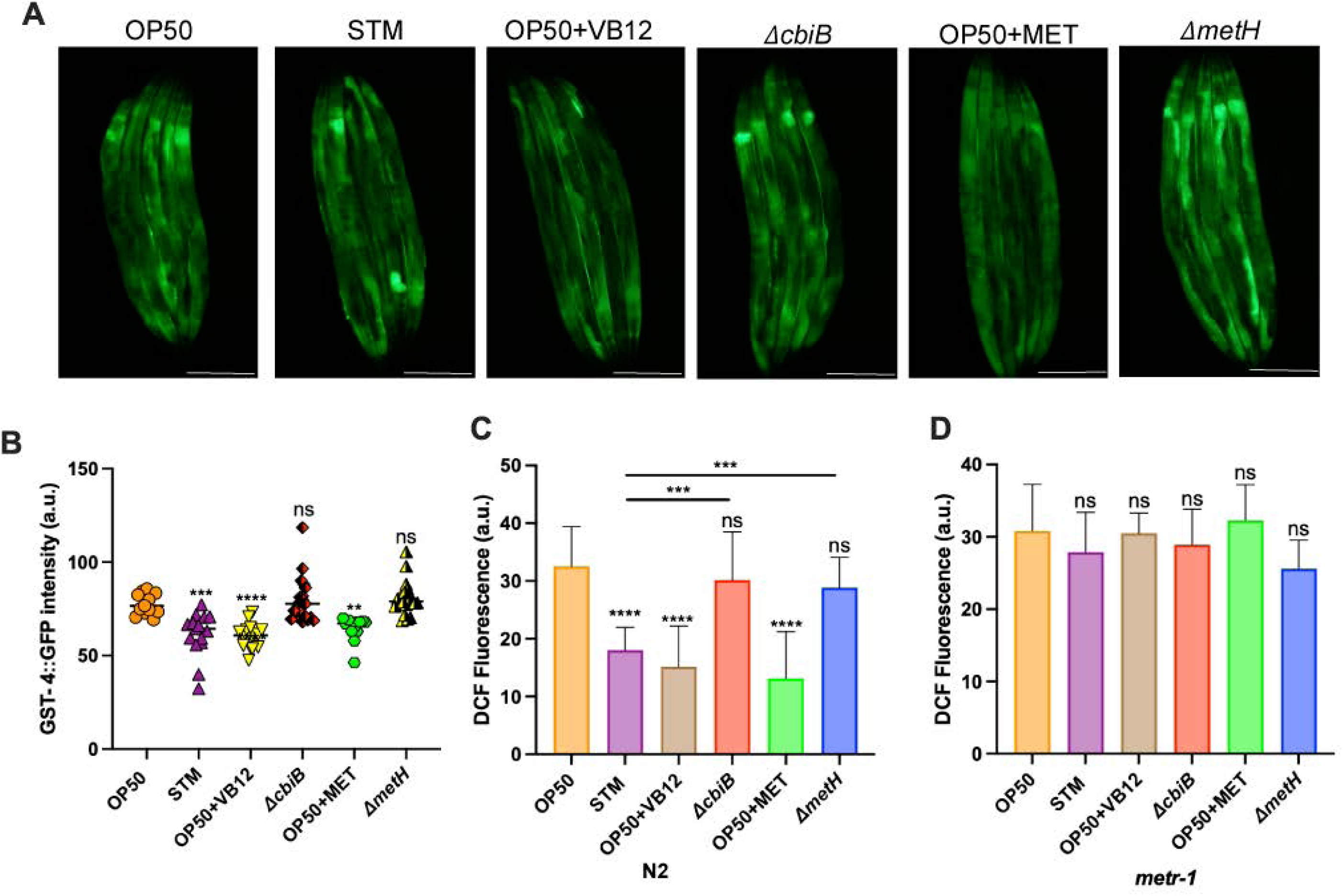
*Salmonella* Typhimurium increases vitellogenin activity and acts as an antioxidant, restoring vitellogenesis and oocyte development. (A) Representative fluorescence images and (B) Quantification of GST-4::GFP expression in day-1 adult worms under the indicated bacterial conditions. Quantification of fluorescence levels of 2′,7′-dichlorofluorescein (DCF) in (C) N2 and (D) *metr-1* animals under the indicated bacterial conditions. An ordinary one-way ANOVA, post-hoc Tukey HSD test was performed. *\*\*\*\*P<0.0001*; *\*\*\*P < 0.001; **P < 0.01; * P < 0.05.* Scale bar = 100 μm.

### *Salmonella* infection decreases mitochondrial activity by reducing its biogenesis

Vitamin B12 decreases the functions of HSP-6, a key component of the mitochondrial unfolded protein response in *C. elegans*, and regulates mitochondrial stress response (82). We aimed to assess mitochondrial health by monitoring the expression of the mitochondrial stress reporter transgene*, hsp-6::GFP*, and measuring mitochondrial genomic DNA copy number. The *hsp-6::GFP* transgenic worms were exposed to the bacterial conditions mentioned above. Under WT-STM, *E. coli* OP50 supplemented with vitamin B12 or methionine, the relative mean fluorescence intensity of HSP-6::GFP was significantly decreased. In contrast, the GFP level was comparable in *E. coli* OP50, Δ*cbiB*, and Δ*metH* fed *C. elegans,* suggesting that vitamin B12-mediated methionine metabolism can maintain nematodes’ proteotoxic stress response (Fig. 7A and B). The mtDNA copy number is a marker of mitochondrial biogenesis, as mtDNA replication is integral to this process (83). To examine the mitochondrial biogenesis rate, we analyzed the mtDNA copy number using the mitochondrial ATP synthase gene (*atp-6*), which is crucial for ATP synthesis. Normalization was performed with the housekeeping nuclear gene *ama-1*. The mtDNA copy number is decreased in WT-STM, *E. coli* OP50 supplemented with vitamin B12 or methionine. At the same time, Δ*cbiB* and Δ*metH* fed worms showed levels comparable to *E. coli* OP50 (Fig.7C). Thus, the decreased mitochondrial gDNA and HSP-6::GFP expression in high vitamin B12 diets suggest that increased vitamin B12 may inhibit mitochondrial biogenesis and stress response activation, reflecting improved mitochondrial health and proteostasis.

**Fig. 7.**
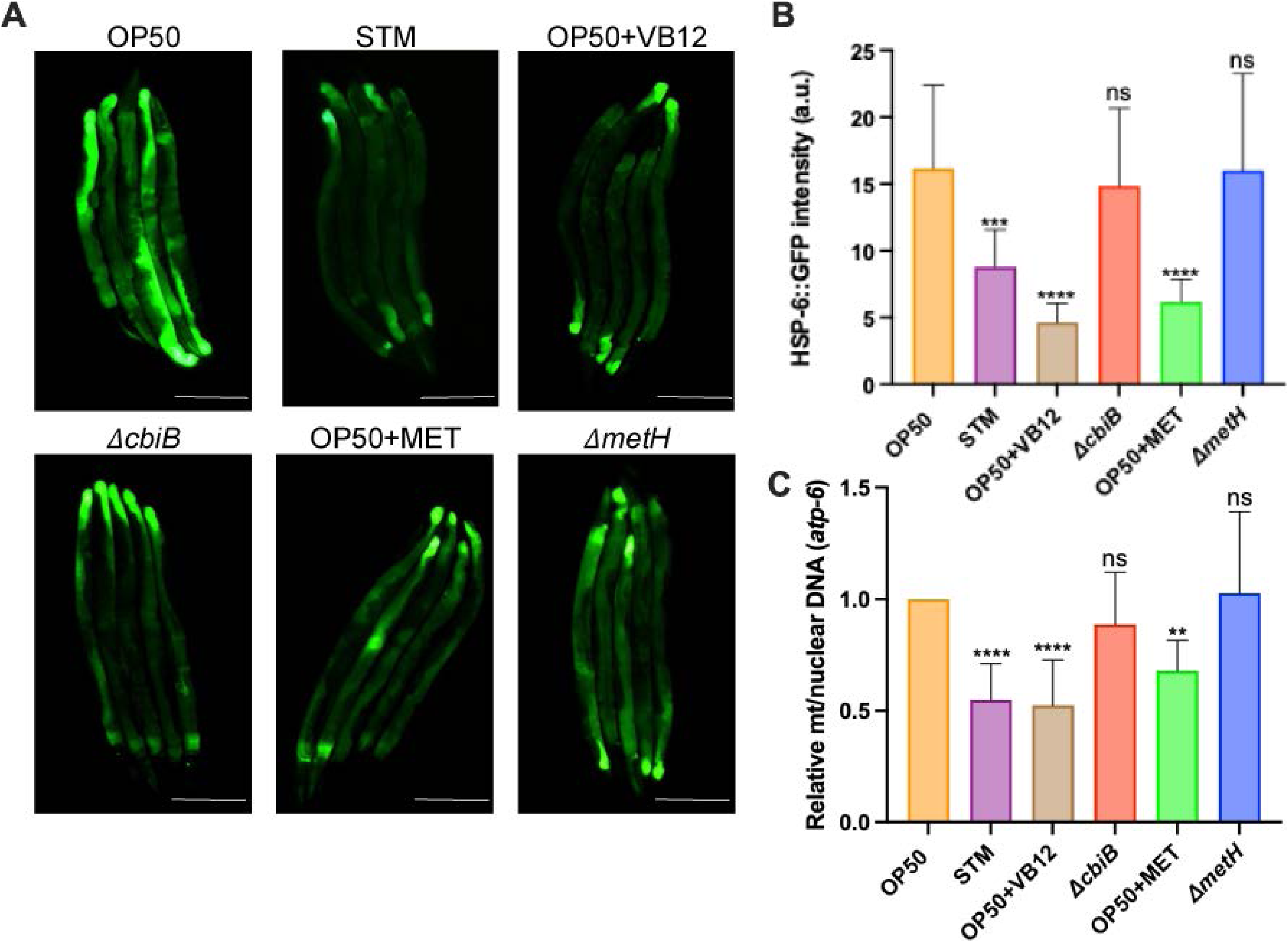
*Salmonella* Typhimurium decreases mitochondrial activity. (A) Representative images and (B) Quantification of GFP levels of *hsp-6::gfp* worms on *E. coli* OP50, wildtype *Salmonella,* Δ*cbiB,* Δ*metH, E. coli* OP50 supplemented with either vitamin B12 or methionine, respectively. HSP-6::GFP expression is reduced in worms grown on WT-STM, *E. coli* OP50 supplemented with vitamin B12 and methionine, respectively. (C) Represents the quantification of total mtDNA copy number from day-1 adult N2 worms on the indicated bacterial conditions by qPCR. WT-STM, *E. coli* OP50 supplemented with vitamin B12 and methionine, respectively, reduces mitochondrial DNA content, while animals fed with *cbiB* and *metH Salmonella* mutants show increased levels comparable to *E. coli* OP50. An ordinary one-way ANOVA, post-hoc Tukey HSD test was performed. *\*\*\*\*P<0.0001*; *\*\*\*P < 0.001; **P < 0.01; * P < 0.05.* Scale bar = 100 μm.

## DISCUSSION

Our previous lab study showed that *Salmonella* infection shortened the lifespan and accelerated developmental progression in *C. elegans* (16). This study demonstrates that *Salmonella* Typhimurium accelerates development and reproductive maturity in *C. elegans* through a vitamin B12-dependent mechanism, rather than via virulence factors (Figs 1 and 2). The relationship between infection and reproduction involves a complex trade-off in which organisms balance immune defense and reproductive effort (84, 85). Understanding immune-reproductive interactions is crucial for developing effective prevention and treatment strategies within the host-pathogen dynamics. *C. elegans* has been used as a model to understand the immunity-reproduction trade-offs and pathogen-specific reproductive strategies (86, 87). Exposure to pathogenic *Pseudomonas aeruginosa* reduces brood size and causes germline shrinkage, whereas exposure to the *gacA* mutant, which is defective in virulence, reverses the germline defect (88). The *Salmonella fepB* enhanced dauer formation and stress resistance in *C. elegans* during infection (40). We also exposed the worms to various avirulent *Salmonella* strains, including SPI-I and SPI-II, as well as non-SPI mutant strains, as listed in Table 1. We observed that these mutants still increased the developmental rate and reproductive maturity, as observed by increased embryo and oocyte production (Figs. 1B and C). Different bacterial diets are a major determinant of *C. elegans* physiology and life-history traits. For instance, a *Comamonas* DA1877 diet accelerates development but reduces fecundity and shortens lifespan compared to the *E. coli* OP50 diet, and these effects are attributed to the nutritional composition rather than pathogenicity (57). Vitamin B12 is vital for methionine synthesis and methylmalonyl-CoA metabolism, making it indispensable for cellular energy production, DNA synthesis, and nervous system maintenance (89, 90). The microbiota’s role is essential to host metabolism, and it provides a framework for understanding the complex interactions among diet, microbiota, metabolism, and disease (21).

*Salmonella hilA* mutant and vitamin B12 supplementation showed comparable progeny and oocyte numbers to those of WT-STM (Fig. 2A-C), indicating that bacterial metabolism, rather than virulence factors, is involved in regulating reproductive development. *Salmonella* exhibits a vitamin B12-rich diet compared to *E. coli* OP50 (Fig. 2D), which contributes to its rapid development in *C. elegans.* Our findings highlight that organisms traditionally classified as pathogens may also benefit their hosts. This duality complicates our understanding of host-microbe interactions and highlights the need for a more detailed approach to microbial ecology. We found that *Salmonella* mutants deficient in cobalamin (*cbiB*) and methionine (*metH*) biosynthesis pathways reduced the developmental acceleration observed in *C. elegans* when fed a vitamin B12-rich diet (Fig. 3A and 3C), indicating a crucial role for these pathways in development. Dietary regulation of cellular metabolism requires adaptations related to vitamin B12 and one-carbon metabolism. Specifically, the folate cycle influences germline development (91), while metabolites of the methionine cycle enhance RAS/MAPK traits (61). The *metr-1* mutant of the worm, defective in vitamin B12-dependent methionine synthesis, did not show accelerated development when exposed to WT-STM and *E. coli* OP50 supplemented with vitamin B12, suggesting that *C. elegans* utilizes bacterial vitamin B12 to enhance its methionine synthesis (Fig. 4B and F) and that this process contributes to its development. In the *metr-1* mutant, the absence of functional methionine synthase disrupts the methionine cycle, preventing the conversion of dietary vitamin B12 into its active forms and resulting in impaired embryo and oocyte production (92). These defects are likely due to impairments in downstream processes, such as SAM production and histone methylation, which are vital for oogenesis (93). Methionine supplementation restored normal embryonic development and oocyte production in the *metr-1* mutant compared to WT-STM (Fig. 4C and G). Vitamin B12 is essential for regulating reproduction in *C. elegans* by enhancing vitellogenin production through the methionine/SAM cycle (31, 82). This process affects gene expression, improves oocyte quality, and optimizes reproductive timing, allowing organisms to adjust their strategies in response to dietary metabolites and the microbiota (61, 94, 95). Methionine supplementation increased antioxidant defense by elevating superoxide dismutase (SOD), catalase (CAT), and glutathione peroxidase (GPx) activity while reducing malondialdehyde levels, thus mitigating oxidative stress during heat exposure (96). *S*. Typhimurium exposure increases vitellogenin production (Fig. 5A-D) and decreases oxidative stress (Fig. 6A-C) in *C. elegans*, likely by improving methionine metabolism. Thus, vitellogenin acts as an antioxidant, protecting organisms from ROS and enhancing resistance to oxidative stress. This protective role has been reported in various species, including nematodes, insects, fish, and vertebrates (97–99).

Mitochondrial abnormalities in oocytes can be inherited, leading to developmental defects and increasing the risks of metabolic disorder in offspring, underscoring the importance of mitochondrial function for transgenerational health (100, 101). We found that mitochondrial function was altered in *S*. Typhimurium-fed worms, with decreased mitochondrial stress responses and biogenesis (Fig. 7A and 7 B). These results highlight the complex metabolic interactions between bacteria and their host (102). *Salmonella*, often seen as a pathogen, provides nutritional benefits due to its high vitamin B12 content, which enhances methionine metabolism in the host and promotes faster development and reproduction. This challenges the view of microbes as merely pathogens, suggesting that *C. elegans* may benefit from *Salmonella* during interaction. The study highlights the implications of vitamin B12 deficiency for other organisms, including humans, and demonstrates that altered mitochondrial function in these worms results in increased mitochondrial stress responses (103, 104). Increased vitellogenin production and reduced oxidative stress underscore the positive effects of bacterial metabolites on reproductive health. Early oocytes exhibit a mitochondrial adaptation that maintains low ROS levels during metabolism, crucial for oocyte quality and developmental potential (105, 106). Vitamin-rich diets led to a reduced number of mitochondria, with a stable membrane potential (107, 108). It also significantly decreased ROS production. These findings suggest that a dietary deficiency in vitamin B12 has a severe impact on mitochondrial health and function. ROS may play a role in regulating mtDNA copy number (109). mtDNA and nuclear genes encode proteins for oxidative phosphorylation. mtDNA is prone to oxidative damage from ROS, leading to mutations linked to aging and mitochondrial disorders (110). Our findings show that exposure to a vitamin B12-rich diet decreases ROS levels (Fig. 6A-C), leading to the inactivation of mitoUPR in worms (Fig. 7A and B), thereby maintaining mitochondrial homeostasis (111).

In conclusion, this study reveals a metabolic interaction between *S*. Typhimurium and *C. elegans* mediated by bacterial vitamin B12, highlighting the role of microbial metabolism in influencing nematode development and reproductive maturity. The full molecular mechanisms underlying *Salmonella’s* effects on vitamin B12 are likely more complex and may require in-depth exploration using proteomics, metabolomics, and other omics approaches.

## MATERIALS AND METHODS

### *C. elegans* culture and maintenance

*C. elegans* N2 (Bristol) strain was used as a wild type (Brenner, 1974) and maintained at 20 °C on standard nematode growth medium (NGM) plates seeded with *Escherichia coli* OP50. Before the experiment, Synchronized L1 larvae were harvested from the gravid *C. elegans* using 5% sodium hypochlorite and 5N NaOH and incubated for 20 hours in an M9 buffer (KH2PO4 0.3%, Na2HPO4 0.6%, NaCl 0.5%, 1 mM MgSO4) with gentle shaking at room temperature. The transgenic worm used in this study: OD95: *ltIs37 [pie-1p::mCherry::his-58 + unc-119(+)] IV. ltIs38 [pie-1p::GFP::PH(PLC1delta1) + unc-119(+)],* VL749: *wwIs24 [acdh-1p::GFP+unc-119(+)]*, CL2166: *dvIs19[(pAF15)gst-4p::GFP::NLS] III* , SJ4100: *zcIs13 [hsp-6p::GFP + lin-15(+)]*, RB755*: metr-1(ok521) II*, RB1434*: mmcm-1(ok1637) III*, VC2428: *sams-1(ok2946),* and DH1033: *bIs1 [vit-2::GFP + rol-6(su10060)] X.* All transgenic worms are maintained at 20 °C.

### Bacterial strains, media, and growth conditions

The following bacterial strains were used: *Escherichia coli* OP50 and *Salmonella* Typhimurium 14028, which served as the wild-type parent strain. *Escherichia coli* OP50 and *Salmonella* Typhimurium were cultured in Luria-Bertani (LB) broth at 37 °C. Δ*hilA*, SPI-1 effector mutants, and Δ*ssaV* SPI-2 mutant*, E. coli* containing pKD4 strain, were grown in LB broth with kanamycin (50µg/ml) at 37 °C, and Δ*phoP* was *grown* in LB broth with chloramphenicol (50µg/ml) at 37 °C*. S*. Typhimurium carrying pKD46 was cultured in LB broth with ampicillin (50µg/ml) at 30 °C. Δ*cobB,* Δ*cobC,* Δ*cobD,* Δ*cobS,* Δ*cbiB,* Δ*pabA,* Δ*metC,* Δ*metE,* and Δ*metH Salmonella* mutants generated in the laboratory, as described below, were used for this study. These mutants were grown in LB broth with kanamycin (50 µg/ml) at 37 °C.

### Generation of cobalamin, folate, and methionine biosynthesis mutant strains of *Salmonella* Typhimurium

*S.* Typhimurium mutant strains were generated using a one-step gene inactivation process (112). Briefly, the *Salmonella* carrying the red helper plasmid pKD46 was grown overnight in LB medium with ampicillin at 30 °C. This overnight culture was subcultured at a 1:100 ratio in Super Optimal Broth (SOB) medium supplemented with 1 M MgSO4, 1 M MgCl2, and 1 M arabinose, and then incubated until the optical density (O.D.) reached 0.4-0.6. Electrocompetent cells were prepared by washing the culture with ice-cold 10% glycerol. The kanamycin cassette from the pKD4 plasmid was amplified using cassette-specific primers mentioned in Table 2 for the genes of interest (*cobB, cobC, cobD, cobS, cbiB, pabA, metC, metE,* and *metH*) and purified. The PCR product was transformed into the pKD46-*Salmonella* strain by electroporation, then a pre-warmed Super Optimal Broth with Catabolite repression (SOC) medium was added and incubated at 30 °C overnight. The culture was then plated on LB agar containing kanamycin (50 µg/ml) at 37 °C for 12 hours, and kanamycin-resistant mutants were confirmed by colony PCR.

**Table 2.**
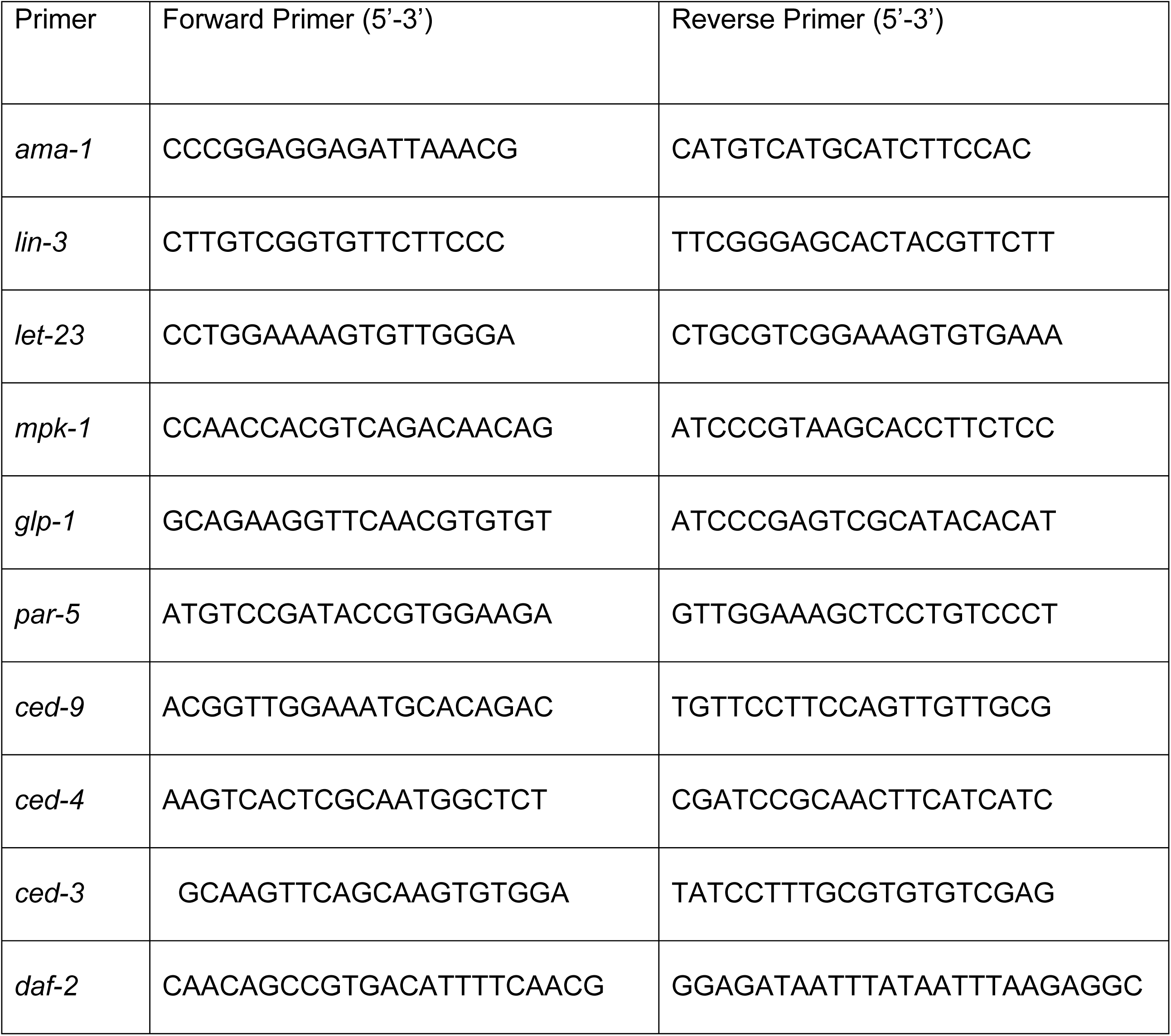
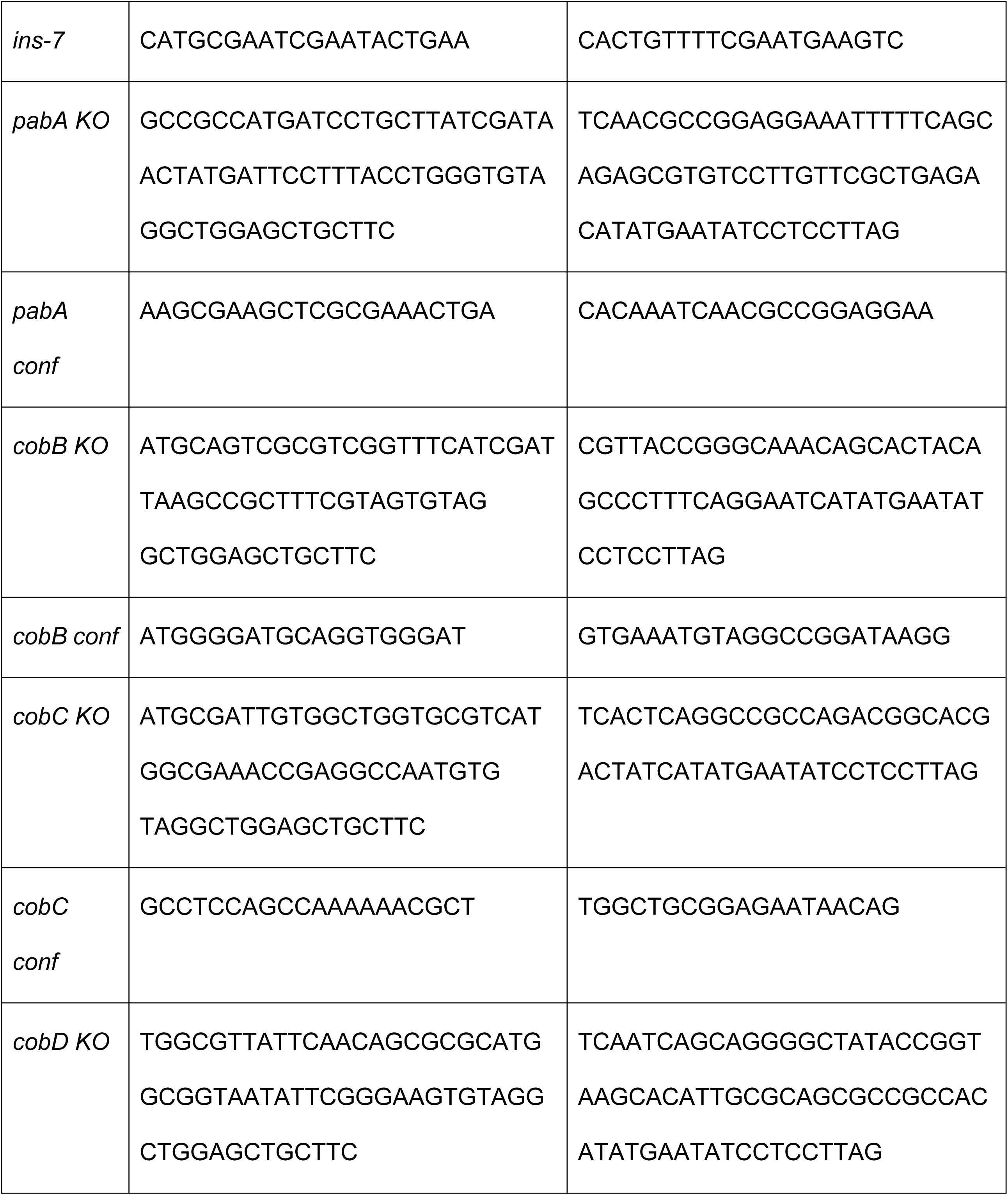
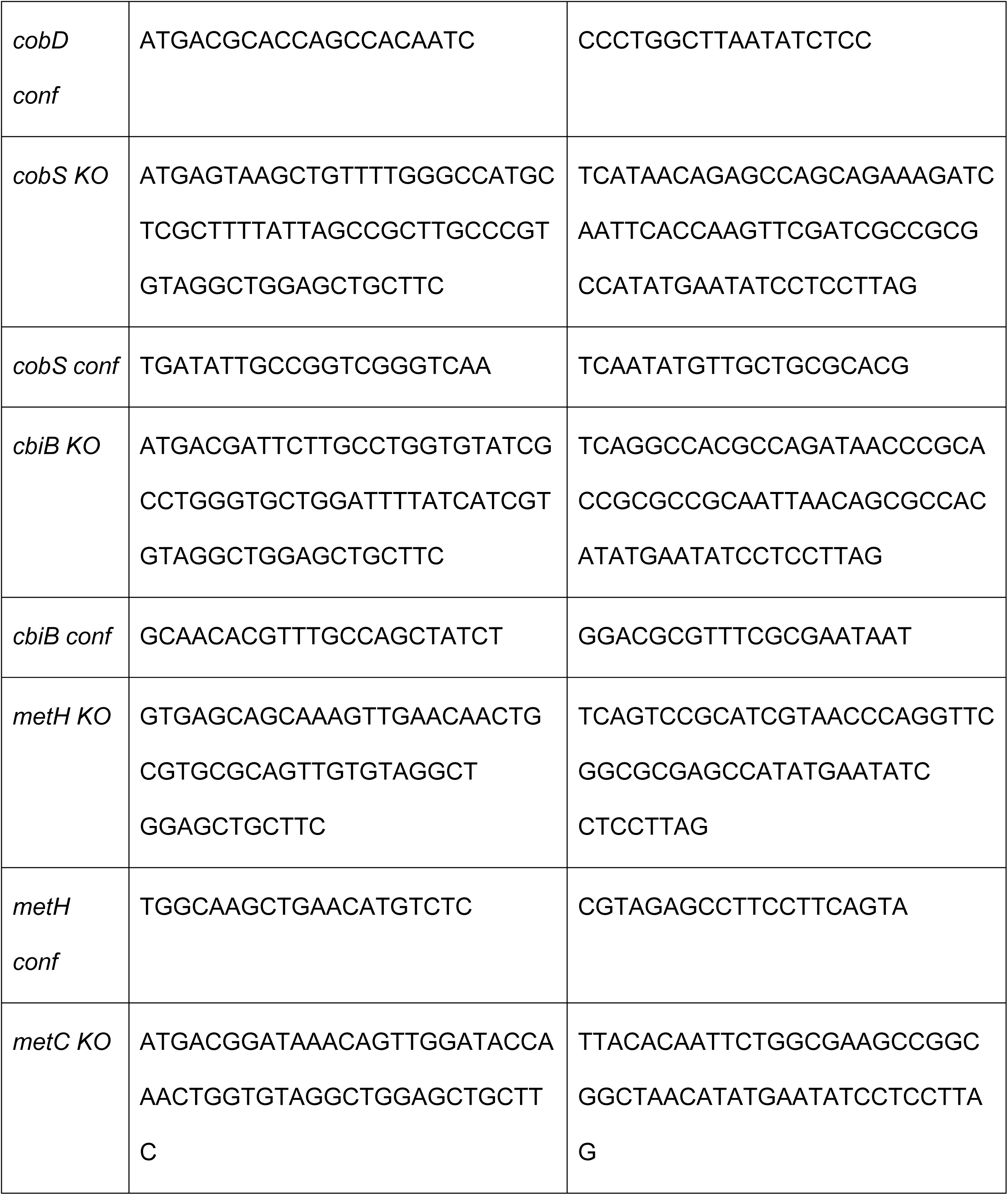
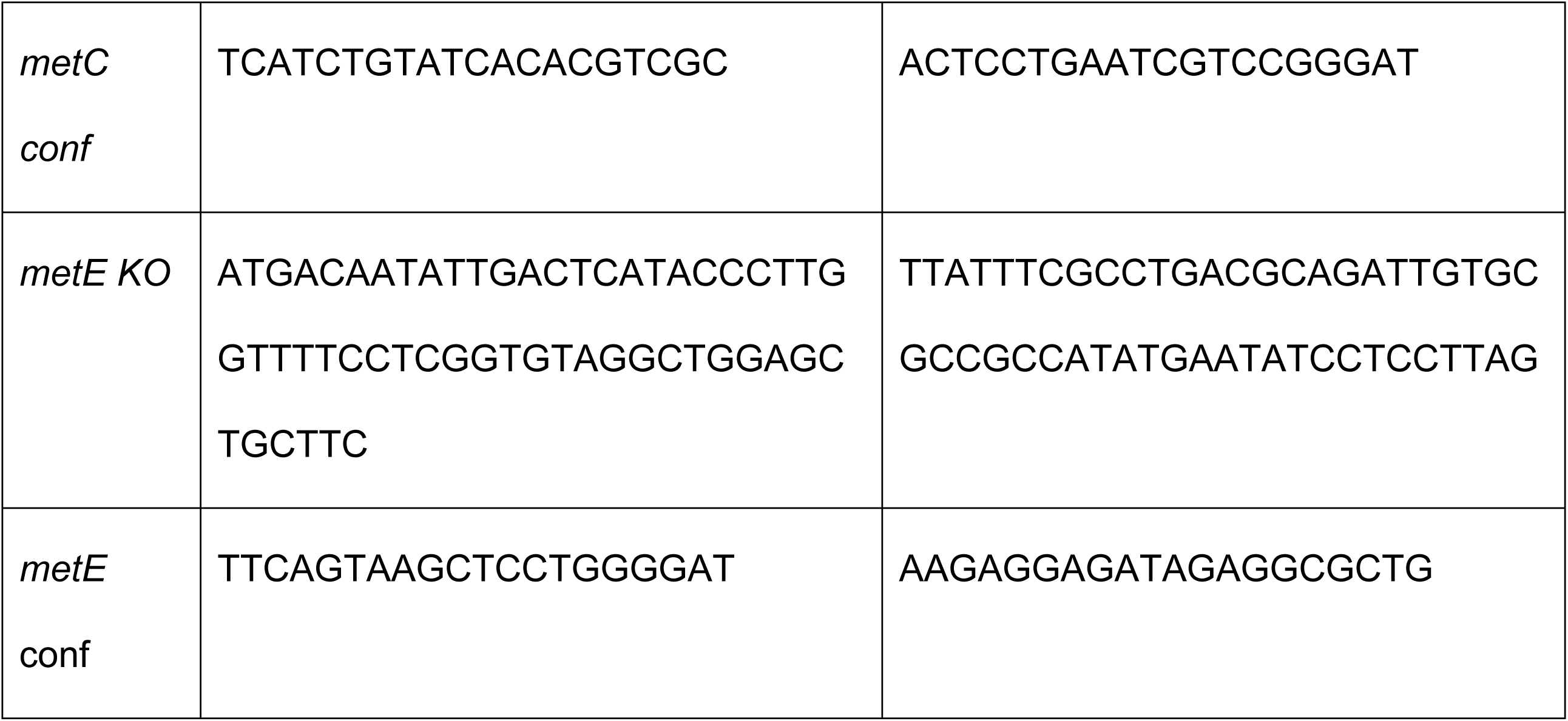
List of primers used in the study

### Exposure of *C. elegans* to *Salmonella* and its mutant strains

Synchronized L1-stage larvae were exposed to wild-type *Salmonella* (WT-STM) and its mutants. The worms were then allowed to grow for 56 hours until they reached young adulthood, and embryo production was quantified. Day-1 adulthood was used to quantify oocyte production.

### Brood size analysis

Brood size of *C. elegans* was determined as described previously (113), with slight modifications by placing a single late L4-hermaphrodite in a freshly seeded plate with *E. coli* OP50, WT-STM, Δ*hilA*, and *E. coli* OP50 supplemented with vitamin B12, allowing it to lay eggs for 24 hours, and transferring it to a fresh plate afterward. This was repeated every 24 hours until the animal stopped laying eggs. Viable progenies (L4 larvae) were counted in each plate after 48 hours and scored. Brood size was expressed as the total number of viable progeny per animal. For each bacterial condition, five hermaphrodite worms were transferred to separate plates, each with one worm.

### Analysis of embryos and oocyte numbers

N2, *metr-1*, *mmcm-1*, and *sams-1* worms were exposed to the bacterial conditions mentioned above for embryo number quantification. Embryos were numbered and scored in 56-hour-old young adult animals in N2, *metr-1*, and *mmcm-1*, while at 72 hours in *sams-1* worms. The oocyte counts were quantified on day 1 adult hermaphrodites of N2, *metr-1*, and *mmcm-1* at 96 hours post-L1-stage in *sams-1* worms. At the indicated time points, *C. elegans* were washed three times in M9 buffer to remove excess bacteria, anesthetized with 25 mM Levamisole in M9 buffer, mounted on 3% agarose pads on glass slides, covered with coverslips, and subjected to a 40X compound microscope. For oocyte quantification using the OD95 strain, the samples were prepared similarly to those of N2 worms and imaged with a 60X objective lens on a Zeiss laser-scanning confocal microscope (LSM 980).

### Vitamin B12 and methionine supplementation

Vitamin B12 (cobalamin) was purchased in its precursor form, cyanocob(III)alamin (CNCbl) (Sigma V2876). Vitamin B12 was dissolved in sterile Milli Q water at a 1 mM stock concentration and added at a final concentration of 64 nM in the NGM plates seeded with the indicated bacterial conditions. Methionine (50 mg/mL) stock was prepared in sterile Milli-Q water and added to NGM seeded with the indicated bacterial conditions at a final concentration of 10 mM.

### Quantification of ACDH-1::GFP, VIT-2::GFP, GST-4::GFP, and HSP-6::GFP expression

The reporter strains VL749, DH1033, CL2166, and SJ4100 were used to visualize ACDH-1, VIT-2, GST-4, and HSP-6 expression, respectively. They were synchronized as stated above to obtain age-synchronized L1 larvae. Approximately 200 aged-synchronized L1 larvae were transferred to each NGM plate seeded with the indicated bacterial conditions, and were incubated at 20 °C for 72 hours. After that, the worms were prepared for fluorescence imaging.

### Fluorescence imaging

Fluorescence imaging was performed by washing the worms with three gravity steps in M9 buffer to remove residual bacteria. They were anesthetized with 25 mM Levamisole and mounted on slides with 3% agarose pads. The animals were then visualized using a Nikon SMZ18 fluorescence stereomicroscope. Quantification of fluorescence intensity was performed using ImageJ.

### Quantitative real-time (q-RT) PCR

*C. elegans* were grown on different bacterial diets until day-1 adulthood, 72 hours after synchronized L1 larvae. For each replicate, approximately 1,000 worms were washed off the plates with M9 buffer and subjected to three gravity washing steps to remove extracellular bacteria. These were collected in Trizol (Invitrogen) and used for RNA isolation per the manufacturer’s instructions. The isolated RNA was treated with DNase (ThermoFisher Scientific) to remove genomic DNA contamination. cDNA was synthesized from 2μg of RNA using the RevertAid First Strand cDNA Synthesis Kit (ThermoFisher Scientific). qRT-PCR was performed using PowerUp SYBR Green Master Mix (Thermo Fisher Scientific) with *ama-1* as the housekeeping gene. The primer sequences are given in Table 2. The 2^-ΔΔCt^ method was used to calculate the relative gene expression.

### Quantification of cytosolic reactive oxygen species (ROS) levels

ROS levels were measured using 2’,7’-dichlorofluorescein diacetate (DCFHDA) (Sigma-Aldrich #35845) (114). Before the experiments, a fresh 50 μM working solution was prepared from a 100 μM stock solution in M9 buffer. Synchronized L1 larvae were grown on different bacterial diets until day-1 adulthood at 20°C. About 100 worms were transferred to 300 μL of dye (150 μL of 50 μM DCFHDA working solution and 150 μL of M9 buffer), giving a final concentration of 25 μM. The samples were incubated in the dark at room temperature for 3 hours with gentle shaking. Worms were then settled by gravity, the supernatant was removed, and the samples were washed three times with PBS plus 0.01% Triton X-100. DCF fluorescence was imaged using a GFP filter on a fluorescence microscope, and at least ten worms per condition and three biological replicates were analyzed.

### Lifespan assay

For lifespan experiments, approximately 100 age-synchronized L1 larvae were transferred to NGM plates seeded with *E. coli* OP50, Wildtype *Salmonella,* SPI-1 effector mutants, such as Δ*hilA,* Δ*sipA,* Δ*sipD,* Δ*sopA,* and Δ*sopB,* and *ΔcbiB,* and Δ*metH, E. coli* OP50 supplemented with either vitamin B12 or methionine, respectively, and grown up to late L4 larval stages, i.e., 48 hours post-L1 stage. 30-40 late L4 larvae were then transferred to NGM plates supplemented with 50 µg/ml of 5-fluoro-2′-deoxyuridine (FUdR) at 20 °C, seeded with the corresponding bacterial strains. Worms were monitored for mortality after 24 hours and then every other day. Animals that did not respond to touch were scored as dead, with at least 3 biological replicates per condition.

### Mitochondrial DNA quantification

As previously described, mtDNA was quantified by a quantitative real-time PCR-based method (115, 116). Briefly, two-day-old adult hermaphrodites were singly picked into 0.2 mL PCR tubes containing 20 µL worm lysis buffer (50 mM KCl, 10mM Tris, pH 8.4, 25 mM MgCl2, 0.45% Tween 20, 60 µg/mL proteinase K). After freezing at -80 °C for 10 minutes, the tubes were incubated on the worm lysis program in a thermocycler at 60 °C for 1 hour, 95°C for 15 minutes, and then 4 °C for 5 minutes. The 5′-GTTTATGCTGCTGTAGCGTG-3′ and 5′-CTGTTAAAGCAAGTGGACGAG-3′ primer pairs were used for measuring mtDNA levels. These primers hybridize in the *atp-6* gene in the mitochondrial genome (mtDNA). The results were normalized to genomic DNA amplified with the 5′-TGGAACTCTGGAGTCACACC-3′ and 5′-CATCCTCCTTCATTGAACGG-3′ primer pair, which hybridizes to the *ama-1* gene.

Quantitative PCR was performed using PowerUp SYBR Green master mix (Thermo Fisher Scientific) on a Bio-Rad CFX96 real-time PCR system, and the experiment was repeated three times. The 2^-ΔΔCt^ method was used to calculate the relative gene expression.

### Quantification and statistical analysis

Statistical analysis was performed using Prism 10 (GraphPad). All error bars represent the mean ± standard deviation (SD). For comparisons involving more than two samples, an ordinary one-way ANOVA was conducted, followed by Tukey’s multiple comparisons test. In the figures, asterisks (*) signify statistical significance as follows: *\* P < 0.05; **P < 0.01; ***P < 0.001; **** P < 0.0001*, relative to the appropriate controls. The Kaplan-Meier method was used to calculate survival fractions, and statistical significance between survival curves was determined using the log-rank test. All experiments were conducted at least three times.

## ACKNOWLEDGEMENTS

We thank our colleague Dr. Jogender Singh from IISER Mohali for the critical discussions and providing JS-VL749, CL2166, SJ4100, RB755, RB1434, and VC2428 strains of the *C. elegans.* We acknowledge Prof. Dipshikha Chakravortty, Microbiology and Cell Biology Department, IISc. Bangalore, India, for the WT-STM and Δ*phoP* strain and Prof. Michael Hensel, Institut für Klinische Mikrobiologie, Immunologie und Hygiene, Germany, for various SPI-1 effector mutant strains (Δ*avrA*-STM, Δ*sptP*-STM, Δ*sipD*-STM, Δ*sopB*-STM). We further acknowledge IISER Mohali for the intramural funds, extramural grant from SERB (EEQ/2021/000769), the Department of Science and Technology, Govt. of India (FIST grant # SR/FST/LS-II/2017/97 to the Department of Biological Sciences, IISER Mohali), and C.D. is a recipient of UGC, SRF fellowship, and acknowledges the same. The UMN Caenorhabditis Genetics Center (CGC) in Minnesota is acknowledged for supplying the worm strains (OD95 and DH1033) used in the study.

## Author’s contribution

VDN conceived the idea and supervised the study. VDN and CD planned and designed the experiments. CD performed all the experiments, VDN and CD analyzed the data and wrote the manuscript.

## Supplementary figure legends

Fig. S1 Gene expression related to various signaling pathways of *C. elegans* exposed to *E. coli* OP50, WT-STM*, ΔhilA*, *ΔavrA*, *ΔsptP*, *ΔsipD*, *ΔssaV*, *ΔygiM*, *ΔrspA*, *ΔsopB*, *ΔmgtC*, *ΔphoP*, and *ΔfepB.* (A) Genes such as *lin-3, let-23, mpk-1, par-5, glp-*1 are involved in the RAS/MAPK signaling pathway during oogenesis, (B) *daf-2* and *ins-7* are involved in the Insulin/IGF-1 signaling pathway, and (C) *ced-9, ced-4,* and *ced-3* are involved in the programmed-cell death pathway. An ordinary one-way ANOVA, post-hoc Tukey HSD test was performed. *\*\*\*\*P<0.0001*; *\*\*\*P < 0.001; **P < 0.01; * P < 0.05.* (D) Representative Kaplan-Meier plot comparing survival of N2 worms exposed to the indicated bacterial conditions.

Fig. S2 There are no growth defects in the cobalamin, folate, and methionine *Salmonella* synthesis mutants in LB media. (A) Deletion of genes involved in the cobalamin synthesis pathway, such as *cobB, cobC, cobD, cobS,* and *cbiB* genes, does not affect its growth in LB media. The *pabA* gene, responsible for folate synthesis, shows no growth defect. (B). Deletion of genes*, metC, metE*, and *metH,* involved in the methionine synthesis pathway, does not affect bacterial growth in LB media.

Fig. S3 *Salmonella* Typhimurium accelerates development through vitamin B12-dependent metabolism in *C. elegans.* (A) Represents the quantification of embryos and (B) Oocyte production in *mmcm-1* animals on the indicated bacterial conditions. An ordinary one-way ANOVA, post-hoc Tukey HSD test was performed. *\*\*\*\*P<0.0001*; *\*\*\*P < 0.001; **P < 0.01; * P < 0.05.* Fig. S4 (A) Representative fluorescence images and (B) Quantification of VIT-2::GFP expression in *vit-2::gfp* transgenic worms exposed to *E. coli* OP50, wildtype *Salmonella, E. coli* OP50 supplemented with either vitamin B12 or methionine, respectively, for 24 hours from L4 onwards. GFP levels were increased in the day-1 worms exposed to *E. coli* OP50 supplemented with vitamin B12 and methionine, respectively. (C) Representative fluorescence images and (D) Quantification of GST-4::GFP expression in *gst-4::gfp* transgenic worms exposed to *E. coli* OP50, wildtype *Salmonella, E. coli* OP50 supplemented with either vitamin B12 or methionine, respectively, for 24 hours from L4 onwards. The worms exposed to *E. coli* OP50 supplemented with vitamin B12 and methionine, respectively, show decreased GFP expression compared to *E. coli* OP50, while those exposed to STM show GFP expression comparable to *E. coli* OP50. An ordinary one-way ANOVA, post-hoc Tukey HSD test was performed. *\*\*\*\*P<0.0001*; *\*\*\*P < 0.001; **P < 0.01; * P < 0.05.* (E) Representative Kaplan-Meier plot comparing survival of N2 worms exposed to *E. coli* OP50, wildtype *Salmonella,* Δ*cbiB,* Δ*metH, E. coli* OP50 supplemented with either vitamin B12 or methionine, respectively. Scale bar = 100 μm.

